# Small molecule inhibition of the mitochondrial lipid transfer protein STARD7 attenuates influenza viral replication

**DOI:** 10.64898/2026.01.20.700505

**Authors:** Shipra Sharma, Jihan Adonis, Shaochen You, Kevin Hartenbower, Vedita A Singh, Dale Allen, Oyahida Khatun, Saikat De, Rachel Sattler, Jonathan A. Covel, Kris M White, Laura Martin-Sancho, Ayush Mehta, Steven H. Olson, Naoko Matsunaga, Adolfo García-Sastre, Michael J Bollong, Megan L Shaw, Nisar Farhat, Sumit K Chanda

## Abstract

The increasing appearance of drug-resistant and zoonotic influenza strains highlights an urgent need for host-directed antivirals that offer broad-spectrum activity and a higher barrier to resistance. Here, we describe the characterization of M4, a small-molecule identified from a high-thoughput screen that potently inhibits influenza A and B viruses. Mechanistic studies reveal that M4 suppresses replication by causing nuclear retention of the viral ribonucleoprotein (vRNP) complex. Chemoproteomic profiling identified the lipid transfer protein STARD7 as the primary cellular target, and genetic depletion of STARD7 phenocopies the antiviral effects of M4. Additional studies localizes the M4 binding site to cysteine 302 within the lipid-binding domain of STARD7, supporting a model in which STARD7-dependent lipid transfer activity promotes efficient vRNP trafficking and nuclear export. Combining M4 with baloxavir enhances antiviral efficacy in a murine infection model, providing in vivo support for a host-directed strategy. Together, these results identify STARD7 as a metabolic checkpoint licensing vRNP export and establish a proof of concept for therapeutic intervention with small molecule inhibitors.

**Author Summary:** Influenza viruses continue to cause widespread illness and pose an ongoing pandemic threat, in part because existing antiviral drugs can lose effectiveness as the virus evolves resistance. To address this challenge, we focused on identifying therapies that target host cell processes required for viral replication, rather than viral proteins themselves. In this study, we describe a small molecule, M4, that inhibits replication of both influenza A and B viruses by blocking a critical step in the viral life cycle.

We found that M4 acts on a host protein called STARD7, which helps move certain lipids to the right places inside cells. When STARD7 is inhibited, the influenza viral ribonucleoprotein (vRNP) complex becomes trapped in the cell nucleus and cannot reach the cytoplasm, preventing the virus from completing its replication cycle. Disrupting STARD7 genetically produces the same effect, confirming that this host protein is important for influenza replication. These findings point to a previously unrecognized host gating mechanism that controls nuclear export of the vRNP complex during infection.

Although M4 alone showed limited activity in animals, combining it with an existing antiviral drug strongly improved antiviral efficacy. Together, our results reveal a new host pathway that influenza viruses rely on and support host-directed combination approaches to strengthen antiviral treatment and help counter drug resistance.

## INTRODUCTION

Influenza A viruses (IAV) are recognized for causing seasonal epidemics and occasional pandemics (Krammer et al., 2018). These events result in significant morbidity and mortality, particularly among vulnerable populations, often straining healthcare resources (Fischer et al., 2014). Influenza A virus subtypes H1, H2, and H3 have caused four historic pandemics, and recent outbreaks involving the H5, H7, and H9 IAV subtypes indicate their potential for zoonotic transmission and associated pandemic risk (Harrington et al. 2021). Of particular concern is avian H5N1 IAV that is now endemic among wild birds globally, resulting in multiple outbreaks in poultry and most recently also in dairy cows (Webby and Uyeki 2024). Although human infections have been rare thus far, and no sustained human-to-human transmission has been reported, the high number of H5N1 viruses circulating in nature and their proved ability to evolve to transmit in some mammals (marine mammals, minks and cows) raise increased concerns on their human pandemic potential.

Regardless of strain, the standard of care treatments for influenza infection are direct-acting antivirals. The neuraminidase (NA) inhibitors-oseltamivir (Tamiflu), zanamivir and peramivir- are approved for the treatment of influenza A and B and act by preventing efficient release of virions from infected cells (McKimm-Breschkin, 2013). Baloxavir marboxil targets the polymerase acidic (PA) endonuclease subunit of the influenza virus polymerase complex, thereby disrupting viral RNA transcription and replication (O’Hanlon & Shaw, 2019). Favipiravir is a nucleoside analog inhibitor of viral RNA-dependent RNA polymerases, suppressing replication of viral genomes, but is currently only approved in Japan for the treatment of pandemic influenza, not seasonal influenza (Furuta et al., 2013). All of these drugs have been clinically demonstrated to reduce the duration of influenza symptoms and may also limit the spread of the virus to others (Hayden et al. 2018; Ison et al. 2013; Hayden et al. 1999; Treanor et al. 2000). However, their efficacy can be compromised by the rapid emergence of resistant viral variants. In fact, M2 inhibitors (i.e. amantadine, rimantidine) are no longer clinically useful as all human circulating influenza A viruses are resistant (Leonov et al., 2011). The emergence and rapid spread of oseltamivir resistant H1N1 influenza A viruses were cause for concern in 2008, but these viruses were replaced by the 2009 H1N1 pandemic virus that is oseltamivir sensitive (Hurt et al. 2009). Yet, sporadic cases of oseltamivir resistance continue to be identified, including the NA-S247N mutation in 3 human cases of influenza H5N1, which has been reported to reduce susceptibility to oseltamivir (Leung et al. 2024). Taken together, these observations show that oseltamivir-resistant viruses can emerge sporadically and persist in circulation. Lastly, baloxavir marboxil is also sensitive to the emergence of resistance mutations, as observed during clinical trials (Omoto et al., 2018). To address therapeutic challenges for both seasonal influenza and emerging pandemic influenza strains, there is an urgent need for the development of novel antiviral agents, particularly with broad-spectrum activity and a higher barrier to resistance.

Influenza virus replication relies on a significant compendium of host (cellular) proteins to complete an infectious replication cycle, many of which have been comprehensively elucidated though several OMICs-level studies (König et al. 2010; Watanabe et al. 2014; Tripathi et al. 2015; Wang et al. 2017; Haas et al. 2023). Host-directed therapies against viruses represent a growing area of interest in antiviral research and treatment strategies. Targeting host proteins can reduce the likelihood of the virus developing resistance but may also increase the risk of side effects due to disruption of normal cellular functions (Kumar et al., 2020). To date, few such therapies have been developed. Most host-directed drugs used in the clinic have immunomodulatory properties that boost natural antiviral responses (e.g. interferon), or that dampen immunopathogenesis due to viral infection (e.g. corticosteroids) (Badia et al., 2022; Kumar et al., 2020; Tanaka et al., 2022).

We have previously reported a high-throughput screen assessing the activities of approximately 1 million compounds in a cell-based assay of influenza virus replication (White et al. 2015). In the present study, we characterize one of the active antiviral compounds, M4, which possesses broad-spectrum antiviral activity against several influenza virus subtypes both ex vivo, and in combination with DAAs, in vivo. This, combined with the inability to select for resistance, suggested a host-directed mechanism of antiviral activity. Employing a chemoproteomics approach, we determined that M4 targets STARD7 (StAR-related lipid transfer domain containing 7). We find that pharmacological and genetic inhibition of STARD7 abrogates nuclear export of influenza vRNPs without inhibition of CRM-1 dependent nuclear export

Although the XPO1/CRM1 pathway is known to mediate nuclear export of this complex, it remains unclear how host lipid metabolism, intracellular trafficking machinery, and signaling pathways converge to license formation of the vRNP complex. vRNP nuclear export is essential for productive infection and requires assembly of nuclear vRNPs with the viral proteins M1 and NEP to form an export-competent complex. Notably, influenza hemagglutinin signaling has been shown to activate host kinase cascades, including the Raf–MEK–ERK–RSK axis, which promotes phosphorylation of NP and facilitates its interaction with M1, thereby licensing nuclear export of vRNPs. How these signaling events are governed by cellular processes, including cellular metabolic state, membrane composition, or lipid-transfer pathways, to regulate the IAV nuclear export checkpoint, remains poorly defined. A more mechanistic understanding of how viral cues and host metabolic and signaling pathways intersect to control vRNP trafficking may reveal new opportunities for therapeutic intervention to block influenza virus replication.

Our findings implicate STARD7, a regulator of phospholipid trafficking, as a critical host factor required for efficient vRNP nuclear export. M4 engages STARD7 within its lipid-binding pocket, a region essential for phosphatidylcholine transfer and mitochondrial lipid homeostasis, linking disruption of this activity to impaired influenza replication. Together, these findings position STARD7 as a metabolic gatekeeper of vRNP nuclear egress and highlight lipid-handling pathways as tractable targets for host-directed antiviral development.

## RESULTS

### Antiviral activity of M4 against influenza viruses

We have previously reported a high-throughput screen of approximately 1 million small molecules using a recombinant influenza A/WSN/33 virus expressing Renilla luciferase, enabling the capture of inhibitors across all stages of the viral life cycle (White et al., 2015b). One of the most potent compounds identified through this analysis was a small molecule designated as M4 (structure shown in Figure 1A), which exhibited a half-maximal inhibitory concentration (IC50) <0.6 μM, and a selectivity index (SI) >10 against influenza A/WSN/33 (H1N1) virus (WSN) in the human lung epithelial cell line (A549), mouse embryonic fibroblast cells (MEF), and primary human tracheal bronchial epithelial (HTBE) (Figure 1B-D; Table 1). The antiviral activity of M4 against different influenza A virus subtypes and influenza B virus was further confirmed using a multicycle growth assay and plaque assay readout (Figure 1E, Table 1). M4 potently inhibited influenza B/Yamagata/16/88 virus with an IC_50_ of 0.03 μM, and an attenuated highly pathogenic avian influenza H5N1 strain (A/Vietnam/1203/04 HALo), with an IC_50_ of 0.09 μM. It was also potent against influenza A/Wyoming/03/03 (H3N2) and influenza A/California/04/2009 (H1N1) viruses, yielding IC_50_s of 0.12 μM and 0.2 μM, respectively (Figure 1E). Overall, treatment with 1 μM M4 resulted in a >1.5-log10 reduction in infectious viral particle release by cells infected for 24 hours at MOI of 0.1 with influenza A or influenza B virus (Figure 1E).

**Figure 1.**
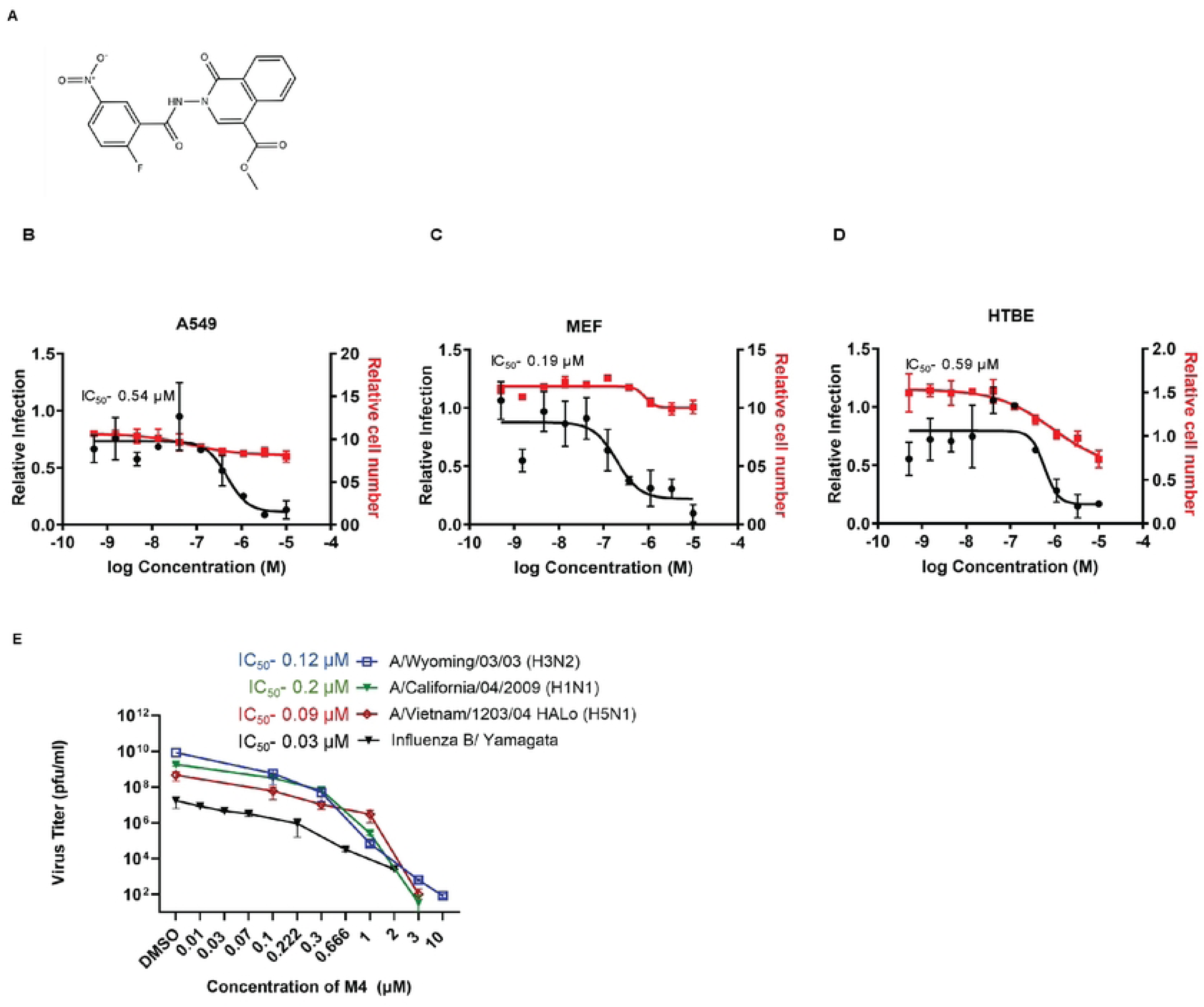
**The M4 compound is a broadly-acting influenza virus inhibitor: (**A) Chemical structure of M4. (B) After pretreatment with M4 for 16h at the indicated concentrations, A549 cells were infected with influenza A/WSN/33 virus at MOI of 0.1, (C) MEF cells were infected with at MOI of 0.5 and (D) HTBE cells were infected at MOI of 0.25. 48h post infection, cells were fixed, stained for NP, and analyzed with a high content immunofluorescence imager. Percent infection was calculated as the ratio of anti-NP-stained cells to DAPI stained cells. Data are normalized by the mean for DMSO-treated wells and are shown as means ± SEM from three independent experiments. Dose-response curves for infectivity (black) and cell number (red) are shown. (E) MDCK cells were pre-treated for 16hs with different concentrations of M4 followed by infection with the indicated influenza A subtypes at MOI of 0.1 or influenza B/Yamagata at MOI of 0.2. Supernatants were analyzed at 24h post infection by plaque assay and data are represented as means ± s.d. of three independent experiments. The IC_50_ values are indicated in Table 1.

**Table 1.**
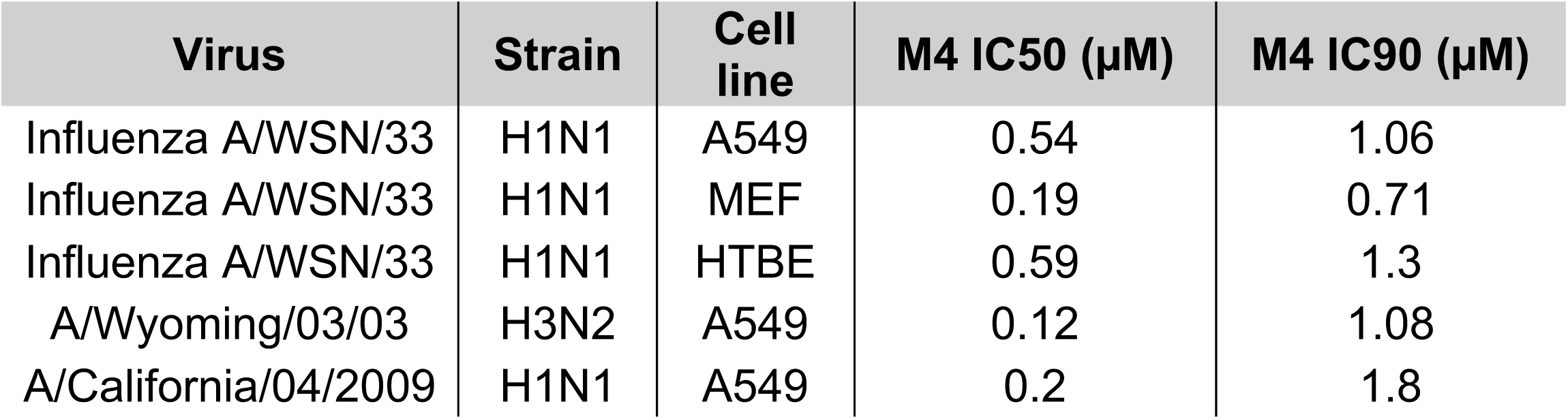
M4 potently inhibits influenza A and B replication in different cell lines.

### M4 inhibits nuclear export of the vRNP complex

To ascertain the stage of the viral life cycle inhibited by M4, we performed a time of addition assay. M4 (at >IC_90_ concentration) inhibited replication of WSN by 2 logs or more when added between 0 and 6 hours post-infection (Figure 2A). This indicates no effect on virus entry as inhibitory activity waned after the 6-hour time-point; it may suggest that M4 acts at post-entry stage in the replication cycle. Using HA/NA pseudotyped vesicular stomatitis virus particles, we further confirmed that M4 does not inhibit viral entry (Figure S1A).

**Figure 2.**
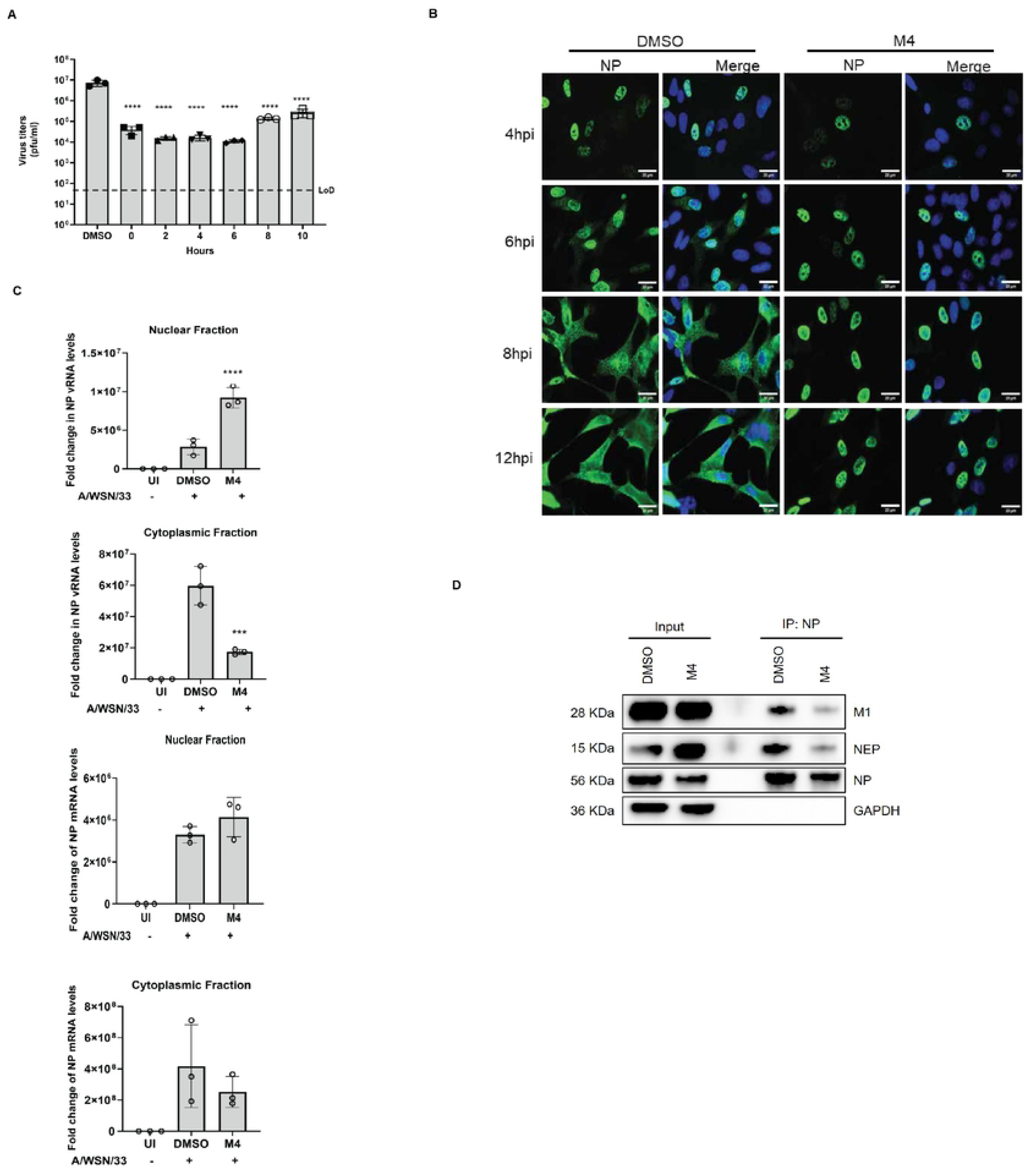
M4 inhibits post entry steps and causes nuclear accumulation of the viral RNP complex: (A) Time-of-addition assay. A549 cells were treated with 3 µM M4 at the indicated times post infection with WSN at MOI of 0.1. Titers were quantified by plaque assay at 24 h post infection (B) Subcellular localisation of NP in M4 treated cells. MDCK cells were treated with DMSO or M4 (3 µM) and infected with influenza A/WSN/33 (MOI 3). Cells were immunostained with anti-NP antibody at 4, 6, 8, and 12hpi, and nuclei visualised with DAPI. Scale bar = *20 µm*. For quantification, 5 fields of view were randomly selected for each condition and the localisation of NP was recorded as present in the “nucleus” or “nucleus/cytoplasm”. (C) Subcellular localization of vRNA and mRNA. A549 cells were pretreated for 2 h with DMSO or 3 µM M4 followed by infection with influenza virus A/WSN/33 at an MOI of 3 for 8 hs. The cells were then subjected to cytoplasmic-nuclear fractionation, followed by RNA isolation from each fraction. The relative nuclear-cytoplasmic distribution of influenza virus NP vRNA (upper panels) NP mRNA (lower panels), was assessed from both subcellular fractions followed by RT-qPCR analysis. Viral RNA abundance in the cytoplasmic fraction was normalized to GAPDH and for the nuclear fraction, with U6 snRNA. The resulting data, relative to that in mock-infected cells, were graphed. Statistical significance was determined with a one-way ANOVA. \*\*\*\**p<0.0001,* \*\*\**p<0.001.* (D) IP description A549 cells were pre-treated with DMSO or M4 (5 µM) and infected with influenza A/WSN/33 (MOI 3). At 8 h post infection, cell lysates were subjected to immunoprecipitation using an anti-NP antibody. Input and NP-immunoprecipitated fractions were analyzed by SDS–PAGE and immunoblotting for viral proteins M1, NEP, and NP. GAPDH served as a control for nonspecific association. Representative immunoblots are shown.

To investigate a potential post-entry mechanism, we asked whether M4 perturbs intracellular trafficking of viral ribonucleoprotein complexes. We therefore visualized the effect of M4 on trafficking of the viral nucleoprotein (NP). Cells were pretreated with M4 (3 μM), infected with WSN at a high multiplicity, and fixed and stained with an anti-NP antibody at 4, 6, 8, and 12 hours post-infection in order to track the movement of NP during a single round of infection (Figure 2B).

In DMSO treated cells, NP exhibits predominantly nuclear staining at the early timepoints, but becomes increasingly cytoplasmic by 8-12 hours post infection as the vRNP is exported from the nucleus for assembly into virions. By contrast, in M4 treated cells, NP staining remains nuclear, even at the later timepoints. To determine if this effect is specific for NP or extends to other components of the vRNP, immunofluorescence staining of NP, PB1, PB2, and PA was performed at 24 hours post infection in the presence of DMSO or M4 (Figure S2A). Over 90% of cells treated with M4 exhibited exclusively nuclear staining of all vRNP proteins, whereas cells treated with DMSO exhibited a diffuse staining, indicating their presence in both the nucleus and the cytoplasm. Similarly, staining for the viral M1 protein over a single infection cycle in the presence of M4, showed that M1, which associates with the vRNP complex to facilitate nuclear export, also remains trapped in the nucleus (Figure S2B). Finally, we examined the localization of influenza virus vRNA and mRNA in M4-treated cells. A549 cells were pre-treated with M4 followed by infection with WSN for 8 hours, allowing only one cycle of replication. Subsequently, subcellular fractionation was performed, and RNA was isolated from cytoplasmic and nuclear fractions. The quantification of negative-sense NP vRNA and positive-sense NP mRNA, in both the cytoplasm and nucleus, was evaluated using strand-specific reverse transcription quantitative polymerase chain reaction (RT-qPCR). Cells pre-treated with M4 exhibited a three-fold enrichment of nuclear NP vRNA, compared to DMSO-treated cells (Figure 2C). Simultaneously, the relative abundance of cytoplasmic NP vRNA was significantly reduced. Nuclear and cytoplasmic NP mRNA levels remained unaffected in M4 treated cells (Figure 2C). Taken together, these findings indicate that M4 causes nuclear retention of the influenza vRNP complex, with both RNA and protein components failing to reach the cytoplasmic compartment. Notably, RanBP1, a host cellular protein that is exported out of the nucleus through the CRM1 complex (Schreiber et al., 2020) did not display nuclear retention following M4 treatment (Figure S2A). As an additional control, we investigated the cellular distribution of STAT1, which translocates to the nucleus upon interferon (IFN) stimulation, followed by re-export into the cytoplasm (Tolomeo, Cavalli, and Cascio 2022). This nuclear export is CRM1-dependent and therefore leptomycin B (LMB)-sensitive. Using a STAT1-GFP fusion, the movement of STAT1 in response to IFN-β was monitored in live cells (Figure S2C). At 1h post IFN-β stimulation in DMSO, M4, and LMB-treated cells, STAT1-GFP was located in the nucleus. By 5hs post-IFNβ stimulation, STAT1-GFP was located both in the nucleus and cytoplasm of DMSO and M4 treated cells, whereas it was primarily localized in the nuclei of LMB-treated cells. These data show that M4 does not act like LMB, suggesting that M4 likely does not targets the nuclear export of influenza virus vRNPs by directly inhibiting the CRM1-mediated export pathway. Since it is known that CRM1 is required for vRNP (Neumann 2000) our results suggest that M4 specifically targets steps upstream of CRM1-dependent export that is unique to vRNP transit.

To investigate whether M4 interferes with the steps that facilitate vRNP export prior to CRM1-mediated export, we examined the association of NP with the viral proteins M1 and NEP, which coordinate formation of export-competent vRNP complexes. In the presence of M4, NP showed markedly reduced association with both proteins (Figure 2D). We confirmed that the immunoprecipitated NP represented bona fide vRNP-associated NP by detecting copurifying vRNA (Figure S2D). These findings support a model in which M4 reduces the assembly of export-ready vRNP complexes by directly or indirectly diminishing NP interactions with M1 and NEP. This would be consistent with the M4-mediated specific inhibition of vRNP nuclear export as compared to other CRM-1 exported proteins.

### M4 is a host-directed antiviral that targets STARD7

We next sought to determine if M4 is a direct-acting or a host-directed antiviral. A distinguishing feature between these two classes of inhibitors is that RNA viruses usually can rapidly acquire resistance to direct-acting antivirals (Irwin et al., 2016). WSN was serially passaged in A549 cells that were treated with M4 (>IC90 concentration) or with DMSO (as a control). After 10 passages, virus titers did not increase in M4 treated cells, indicating that no resistance had developed (Table S1, Figure S3B). This outcome implies that M4 may act on a host cell component, rather than directly targeting the virus itself.

We hypothesized that M4 may covalently bind its target protein due to the presence of its 2-fluoro-5-nitrobenzyl group, which is known to react with cysteine through an S_N_Ar nucleophilic aromatic substitution-based mechanism (Figure 3A) (Shannon et al., 2014). Accordingly, to identify the putative covalent target of M4, we synthesized M4.67, a derivative of M4 bearing an alkyne moiety for use in affinity tagging and enrichment studies (Figure 3B). We next performed chemical proteomics enrichment studies to identify the proteins covalently labeled by M4.67. HEK293T cells were exposed to 1 µM M4.67 for 1h and lysates were subjected to click chemistry reactions to affix biotin azide to labeled proteins. Streptavidin enrichment in denaturing conditions, coupled to shotgun AP-MS/MS proteomic analysis, identified STARD7 as statistically enriched in the immunoprecipitate (Table S2A, Figure 3C,). M4.67 labeling of STARD7 was also confirmed in wild-type A549 cells, as 20 µM M4.67 treatment for 1h followed by biotin azide click reaction and streptavidin enrichment resulted in robust detection of STARD7 by immunoblotting (Figure 3D).

**Figure 3.**
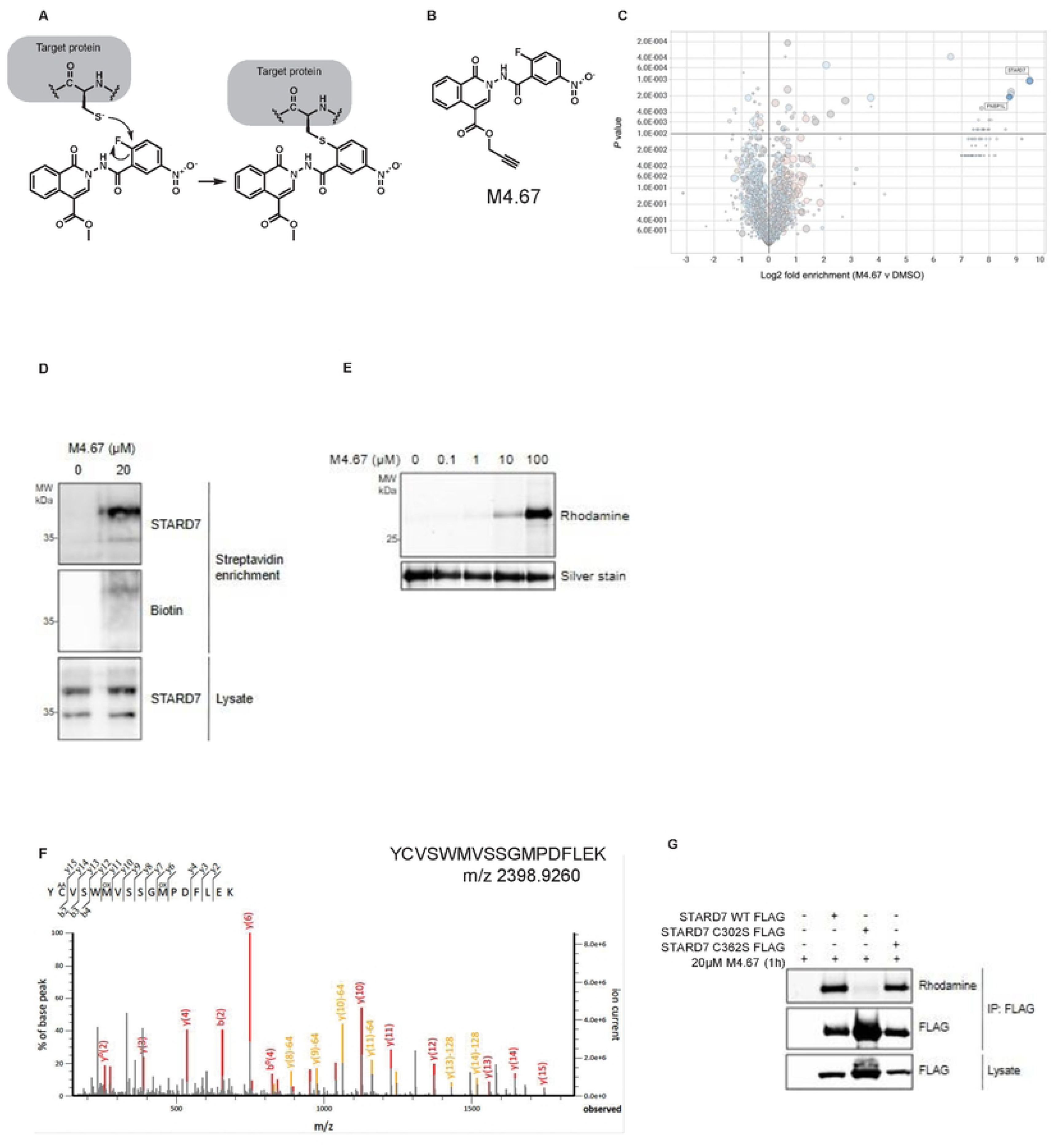
Proteomic enrichment of M4.67 target: (A) Schematic illustrating covalent modification of the target protein by M4 (B) Structure of M4.67 probe (C) Scatter plot of fold enrichment and P value of potential cellular targets from HEK293T cells exposed to M4.67 (1 µM) relative to DMSO. (D) Representative anti-STARD7 and anti-biotin western blot of M4.67 (20 µM) treated A549 cells after streptavidin enrichment of labeled proteins. (E) Representative rhodamine scan and silver stain of M4.67 treated HEK293T cells transfected with STARD7 WT-FLAG transgene after FLAG immunoprecipitation. (F) MS/MS spectra of STARD7 peptide containing modified C302 from anti-FLAG immunoprecipitated material from HEK293T cells expressing STARD7 WT FLAG treated with 20µM M4.67 for 1 hour. (G) Rhodamine fluorescence scan and anti-FLAG Western blot of anti-FLAG immunoprecipitated content of the indicated FLAG tagged STARD7 transgenes overexpressed in HEK293T cells and exposed to M4.67 (20 μM) for 1 hour.

We also found that in HEK293T cells overexpressing FLAG-tagged STARD7, M4.67 dose-dependently labeled STARD7 (100µM to 100nM) after 1h of exposure as determined by rhodamine positivity of anti-FLAG immunoprecipitated material, confirming the results of the chemoproteomic profiling (Figure 3E).

We next sought to characterize the target residue of STARD7 that is covalently labeled by M4.67. We could specifically detect a tryptic peptide fragment adduct containing C302 by MS/MS from HEK23T cells overexpressing STARD7 treated with 20 µM M4.67 for 1 h. (Table S2B, Figure 3F,). Mature processed STARD7 contains two cysteines, C302 and C362 (Flores-Martin et al. 2013). To probe the involvement of these cysteines in the interaction with M4, we generated cysteine mutants of FLAG-tagged STARD7 (C302S and C362S) and overexpressed them in HEK293T cells. After treatment with M4.67, FLAG immunoprecipitation, and click-reaction based conjugation to rhodamine azide, it was found that the C302S mutant no longer bound to M4.67, suggesting that this residue is likely the target of M4.67 (Figure 3G). Cysteine 302 of STARD7 is located within the START domain, which is essential for lipid binding, transport and metabolism (Flores-Martin et al., 2013b). It likely plays a critical role in the lipid-binding function of the protein, either by directly interacting with lipids or by maintaining the structural integrity of the lipid-binding domain.

### STARD7 is required for influenza A virus replication

To determine if STARD7 is a proviral host factor required for influenza A virus replication, gene-specific siRNA knockdown was optimized in A549 cells to achieve a >90% reduction in STARD7 expression (Figure S4A). Infection of STARD7-depleted cells was evaluated by immunostaining of NP and a significant reduction (50%) was consistently observed (Figure 4A). To further validate this observation, virus titers were quantified by plaque assay in STARD7 knockdown cells. Virus release was significantly decreased (>1.5 log_10_) at 24 hours post infection (Figure 4B). Next, we employed CRISPR-mediated genome editing to ablate STARD7 in A549 cells. WSN replication was diminished by approximately 1.5 log_10_ (p<0.005) in these cells, and was comparable to that in M4-treated A549 cells (Figure 4C). Importantly, complementation of knockout cells with wild-type STARD7 re-established viral replication to levels observed in parental A549 cells, while viral replication was not significantly recovered in knockout cells complemented with STARD7 C302S mutant (Figure 4C). Additionally, M4 did not exhibit antiviral activity in STARD7 knockout cells, nor those complemented with mutant C302S compared to wild-type STARD7 and interestingly, supplementing the knockout cells with the full-length STARD7 plasmid significantly restored antiviral activity of M4 (Figure 4D), further indicating that M4 likely exerts its antiviral activity though inhibition of STARD7. Finally, we find that STARD7 knockout, or knockdown, led to retention of vRNP in the nucleus, phenocopying antiviral activities seen with M4 treatment (Figure 4E, S5A). These data strongly support STARD7 as the critical host protein that is targeted by M4 to exert its antiviral activity, underscores the significance of STARD7 as a host factor that is critical for the nuclear export of vRNPs, and provide functional evidence that STARD7 exerts proviral activities that can be pharmacologically targeted to inhibit viral replication.

**Figure 4.**
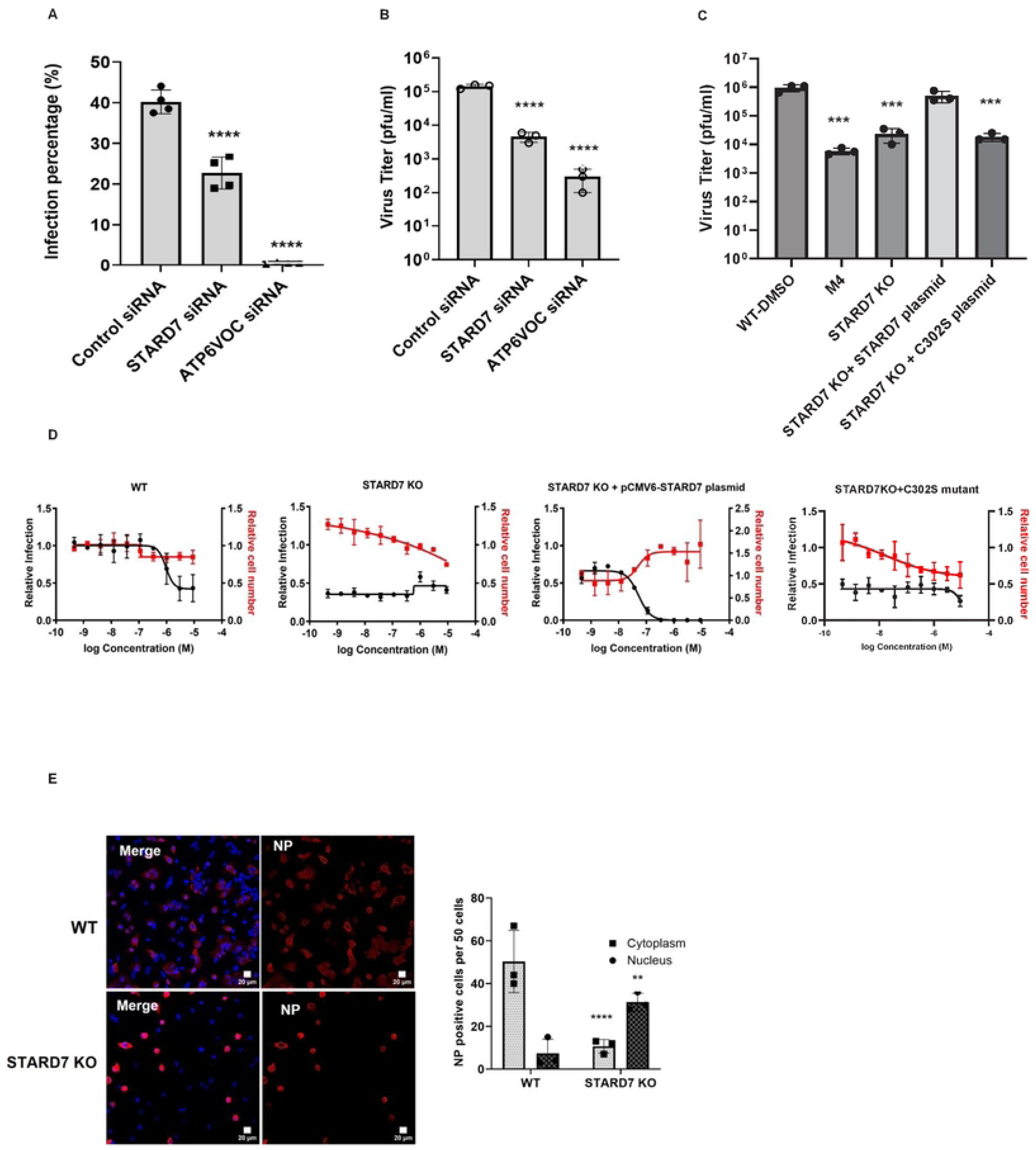
STARD7 is Required for IAV Replication and M4 antiviral activity. (A) A549 cells were transfected with non-targeting control siRNA or siRNA targeting STARD7 or ATP6V0C followed by infection with WSN at MOI of 0.01. At 32 h post infection, the cells were fixed, stained for NP and analyzed with a high content immunofluorescence imager. Percent infection is shown. Statistical significance was determined with a one-way ANOVA. \*\*\*\**p<0.0001.* (B) A549 cells were transfected with non-targeting control siRNA or siRNA targeting STARD7 or ATP6V0C followed by infection with WSN at MOI of 0.01. Titers were quantified by plaque assay at 24 post infection. Statistical significance was determined with a one-way ANOVA. **** p<0.0001. (C) DMSO-treated WT A549 cells , 3 μM M4 treated WT cells, STARD7 KO cells, STARD7 KO + 80 ng of transfected pCMV6-STARD7 and STARD7 KO + 80 ng of transfected C302S plasmids cells were infected with WSN (MOI 0.25) for 48 h followed by quantification of titers by plaque assay. (D) WT A549 cells, STARD7 KO cells, STARD7 KO + transfected pCMV6-STARD7 plasmid and STARD7 KO + transfected pCMV6-STARD7 (C302S) cells were infected with WSN (MOI 0.25) and treated with M4. 48h post infection, cells were fixed, stained and analyzed with a high content immunofluorescence imager. Percent infection was calculated as the ratio of anti-NP-stained cells to DAPI stained cells. Data were normalized by the mean for DMSO-treated wells and represent means ± SEM from thee independent experiments. Dose-response curves for infectivity (black) and cell number (red) are shown. (E) WT and STARD7 KO cells were infected with A/WSN/33 at an MOI of 3 for 16h. The cells were fixed, labelled with anti NP and DAPI. Quantification of NP positive cell per cell where at least fifty cells per condition were quantified for each image with Image J.

### *In vivo* evaluation of M4 in combination with direct-acting antivirals

To evaluate the antiviral efficacy of M4 in vivo, we performed a prophylactic treatment study in a murine model of influenza A virus infection. Mice were administered M4 intraperitoneally at 60 mg/kg twice daily, beginning one day prior to infection with influenza A/WSN/33 virus and continuing through day 3 post-infection (Figure 5A). This dosing regimen was informed by a prior maximum tolerated dose study, in which 60 mg/kg administered twice daily represented the highest dose tested that did not cause significant weight loss (Figure S6B, S6C). As monotherapy, even at near maximally tolerated doses, M4 had minimal impact on viral replication, likely due to insufficient exposure, with lung titers comparable to those observed in vehicle-treated animals, approximately 10⁶ PFU/mL (Figure 5B).

**Figure 5.**
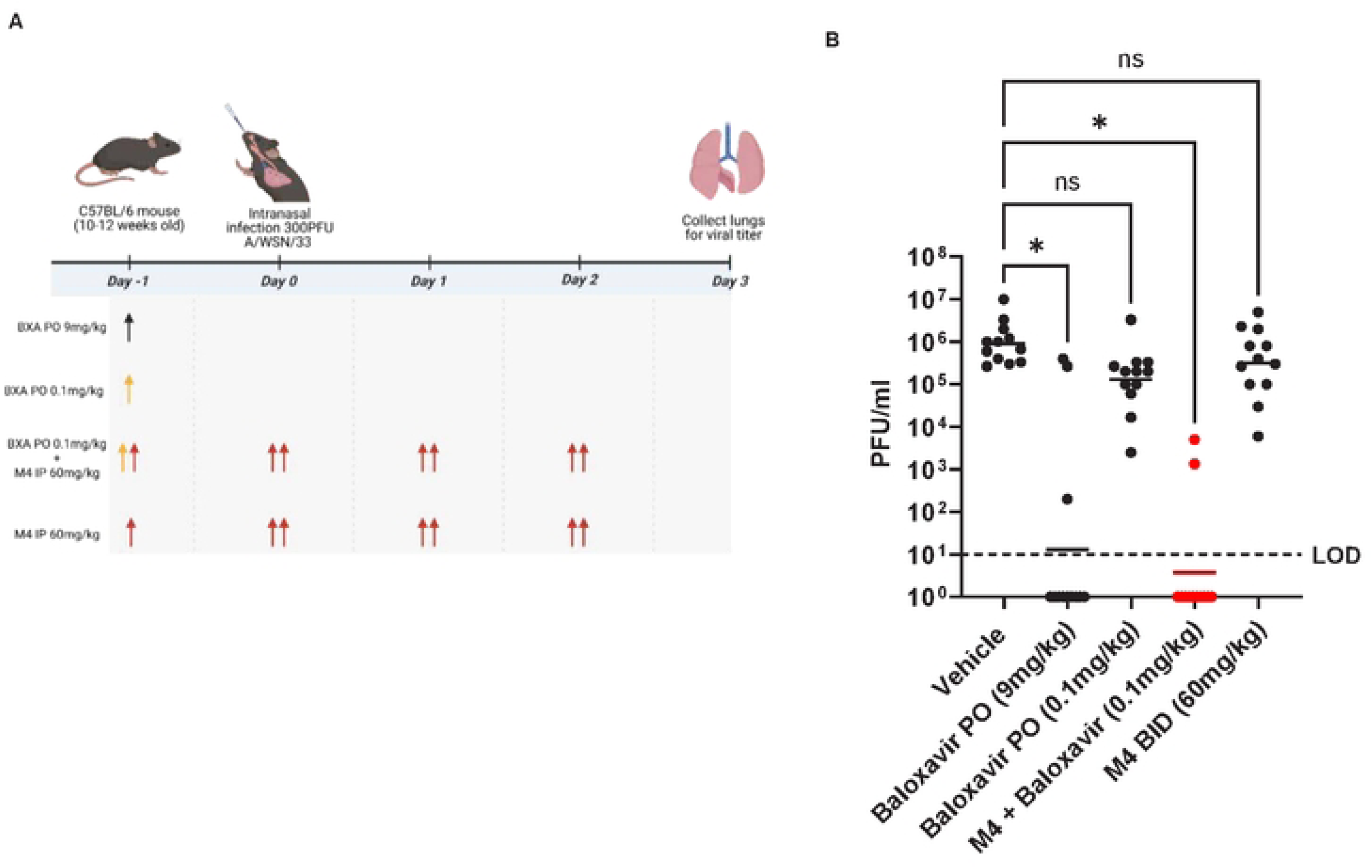
M4 demonstrates in vivo antiviral activity and enhances the efficacy of baloxavir in a mouse model of influenza A virus infection. (A) *Schematic of the experimental design.* C57BL/6 mice (10–12 weeks old) were infected intranasally with 300 PFU of A/WSN/33 (H1N1) on Day 0. Treatments were administered according to the indicated schedule: baloxavir (PO) at 0.1 or 9 mg/kg once daily, M4 (IP) at 60 mg/kg twice daily (BID), or combination therapy with M4 (60 mg/kg IP BID) plus baloxavir (0.1 mg/kg PO). Lungs were harvested on Day 3 post-infection for viral titer quantification. (B) Viral loads were determined by plaque assay and are presented as PFU/mL on a log₁₀ scale. The limit of detection (LOD) is indicated by the dashed line. Statistical significance was assessed by one-way ANOVA with appropriate post-hoc tests; ns = not significant.

Given the limited efficacy of M4 alone, we next evaluated its use in combination with a subtherapeutic dose of a direct-acting antiviral to enhance antiviral activity while limiting drug exposure and reducing the risk of resistance. Baloxavir marboxil was selected for combination studies. Baloxavir was administered prophylactically at 0.1 mg/kg, well below the calculated therapeutic dose of 9 mg/kg, and alone produced only a modest, non-significant reduction in lung viral titers. In contrast, combination treatment with M4 and baloxavir reduced viral titers to below the limit of detection, consistent with an additive or potentially synergistic interaction and supporting further development of this host-directed combination strategy (Figure 5B).

## DISCUSSION

In this study, we identify the small molecule M4 as a potent, covalent inhibitor of influenza virus replication that targets the host factor STARD7. M4 exhibits antiviral activity against a broad range of influenza A viruses, including seasonal H3N2 and H1N1 viruses, and an avian H5N1 subtype virus, which is of major concern to public health due to its potential for zoonotic transmission. M4 also inhibits influenza B virus replication, confirming it as a pan-influenza inhibitor. Further characterization indicates that M4 treatment of influenza virus infected cells traps the viral ribonucleoprotein complex in the nucleus, thereby preventing generation of progeny virions. We find that M4 covalently binds STARD7, and nuclear retention of the vRNP complex is observed in cells depleted of STARD7, which supports the conclusion that this protein is the bona fide antiviral target of M4, and indicates that STARD7 is important for efficient influenza virus replication. Together, these findings define a host-directed mechanism of antiviral activity and reveal a previously unrecognized metabolic checkpoint governing influenza vRNP nuclear export.

STARD7 is a member of the steroidogenic acute regulatory related lipid transfer domain protein family, whose members play critical roles in intracellular lipid transport, particularly involving cholesterol, phospholipids, and other sterols that are essential for membrane composition and signaling (Clark 2012). STARD7 is specifically involved in the transfer of phosphatidylcholine to mitochondrial membranes and is required to maintain mitochondrial membrane integrity and cellular respiration (Horibata and Sugimoto 2010; Rojas et al. 2021; Yang et al. 2017). STARD7 exists in two isoforms: the precursor form STARD7-I, which contains a mitochondrial targeting sequence, and the shorter form STARD7-II, which results from cleavage of STARD7-I by mitochondrial proteases and is localized predominantly in the cytoplasm, plasma membrane, and nucleus (Leman et al. 2009; Angeletti et al. 2008). STARD7-I is responsible for delivery of phosphatidylcholine to mitochondria, while STARD7-II has been implicated in cell migration through ERK1/2 signaling, connexin 43, and integrin β1 pathways (Cruz Del Puerto et al. 2022). The extra-mitochondrial form of STARD7 is also required for transport of coenzyme Q from mitochondria to the plasma membrane (Deshwal et al. 2023). Because phospholipid transfer frequently occurs at mitochondria–ER membrane contact sites, which serve as hubs for lipid exchange and metabolic signaling, STARD7 is well positioned to couple lipid flux and mitochondrial state to downstream cellular processes.

We find several lines of evidence suggesting that M4 exerts its antiviral activity through inhibition of the lipid-transfer function of STARD7. Binding studies indicate that M4 binds within the lipid transferase pocket of STARD7. Mutation of cysteine 302 to serine in this domain abolished M4 binding, and the C302S mutant failed to rescue influenza replication in STARD7-deficient cells, whereas wild-type STARD7 restored replication. These data place the STARD7 lipid-transfer domain, rather than a scaffolding or signaling role alone, at the center of its proviral function. Importantly, M4 treatment phenocopies genetic depletion of STARD7, including nuclear retention of vRNPs, supporting the conclusion that M4 defines a functional checkpoint controlled by STARD7 lipid handling.

Influenza vRNP nuclear export requires assembly of a complex containing NP, M1, and NEP, which subsequently engages the CRM1 export machinery (Neumann 2000; Elton et al. 2001). While CRM1 has been well characterized in this process, the host determinants that license formation of export-competent vRNP complexes remain incompletely understood. Prior work has shown that influenza hemagglutinin–mediated signaling activates host kinase cascades, including the Raf–MEK–ERK–RSK pathway, which promotes phosphorylation of NP and facilitates its interaction with M1, thereby licensing vRNP nuclear export. Critically, we find that M4 inhibits vRNP nuclear export in a manner that does not phenocopy CRM1 inhibition and instead prevents association of NP with M1 and NEP, suggesting that STARD7 functions upstream of CRM1 engagement, and its activity is required for formation of the vRNP complex. In this framework, STARD7-dependent lipid handling emerges as a metabolic checkpoint that integrates upstream viral and host signaling inputs to enable formation of export-competent vRNP complexes.

STARD7 may influence vRNP export indirectly through mitochondrial signaling pathways. Inhibition of the Raf–MEK–ERK cascade results in nuclear retention of vRNPs, a phenotype mediated by the redox-sensitive kinase RSK1, which phosphorylates NP and promotes its interaction with M1 (Pleschka et al. 2001; Droebner et al. 2011; Brunotte et al. 2014). Because STARD7 contributes to mitochondrial lipid handling and cellular redox balance, changes in its activity could modestly influence signaling pathways such as RSK1 that participate in NP licensing. These models are not mutually exclusive and point to an integrated role for lipid metabolism, redox signaling, and viral trafficking in controlling influenza nuclear export. Alternatively, host lipid metabolism is upregulated during influenza infection, and the viral matrix protein M1 directly interacts with phosphatidylcholine, phosphatidylserine, and cholesterol and has been implicated in vRNP nuclear export (Ayari et al. 2020; Baudin et al. 2001). One possibility is that STARD7-dependent phosphatidylcholine trafficking supports the lipid environment required for stable M1–NP–NEP complex assembly, and that inhibition or loss of STARD7 perturbs this lipid-dependent licensing step.

Targeting host pathways required for viral replication offers the advantage of a higher barrier to resistance compared with direct-acting antivirals. Consistent with this, we did not observe resistance to M4 after extended viral passaging. Importantly, M4 does not disrupt CRM1-dependent export of host proteins, suggesting a selective mechanism that may limit toxicity relative to direct CRM1 inhibitors. Although M4 alone showed limited efficacy in vivo at near-maximally tolerated doses, combination with a subtherapeutic dose of baloxavir resulted in complete suppression of viral replication in a murine model. This provides in vivo proof of concept that partial inhibition of a host metabolic checkpoint can strongly potentiate direct-acting antivirals. Such combination strategies may enable dose sparing, improved efficacy, and reduced emergence of resistance.

In summary, this work identifies STARD7 as a metabolic and signaling node that links phospholipid transfer to influenza vRNP nuclear egress. By demonstrating that covalent engagement of the STARD7 lipid-transfer domain blocks vRNP export and viral replication, these findings establish lipid regulation as a previously unrecognized checkpoint in the influenza life cycle and highlight STARD7 as a tractable target for host-directed antiviral discovery.

## MATERIALS AND METHODS

### Cells and viruses

A549 (ATCC CCL-185), MDCK (ATCC CCL-34), and MEF (ATCC CRL-2991) cells were cultured at 37°C in a humidified incubator with 5% CO2 in DMEM (Gibco, Life Technologies) supplemented with 10% FBS (Gibco, Life Technologies), 1 mM sodium pyruvate (Gibco, Life Technologies), 10 mM HEPES (Gibco, Life Technologies), and 100 U/ml of penicillin and 100 μg/ml of streptomycin (Gibco, Life Technologies). HTBE cells (ATCC PCS-300-010) were cultured in commercially available airway epithelial cell basal media following the manufacturer’s protocol (ATCC). A549-doxycycline(dox)-Cas9 cells were generated by transduction with Lenti-dox-Cas9 (Dharmacon) into A549 cells. Following transduction, cells were plated for colony formation and screened for Cas9 expression after doxycycline treatment. Cell viability was assessed via western blotting for STARD7 (PA5-30772, Thermo Fisher) and β-actin (4967, Cell Signaling) and also analyzed using DAPI staining with a high content imager.

Influenza A/WSN/33 (H1N1), A/Wyoming/3/03 (H3N2), A/California/07/2009, A/Vietnam/1203/04-HALo and B/Yamagata/88 viruses were propagated in MDCK cells. A/Vietnam/1203/04 (H5N1) HALo mutant virus is an attenuated H5N1 influenza A virus that lacks the polybasic cleavage site in HA (Steel et al., 2009).

### siRNA knockdown

Adenocarcinoma human alveolar basal epithelial cells, A549, were grown at 37°C in a humidified 5% CO_2_ atmosphere in DMEM supplemented with 10% FBS (Gibco, Thermofisher), penicillin, streptomycin, and amphotericin B mix (PAN-Biotech). A549 cells were transiently transfected with siRNA (40nM siRNA/24-well) using Lipofectamine™ RNAiMAX (Invitrogen, Thermofisher) according to manufacturer’s instructions. All RNA oligonucleotides were synthesized by Dharmacon™ Reagents (Lafayette, CO, USA). Forty-eight hours post-transfection, cells were infected with influenza A/WSN/33 (WSN) at MOI 0.01 (Figure 4A, B) or 6 (Figure 4E, S5A) for 1 hour at 4°C.

siRNA sequences used were:

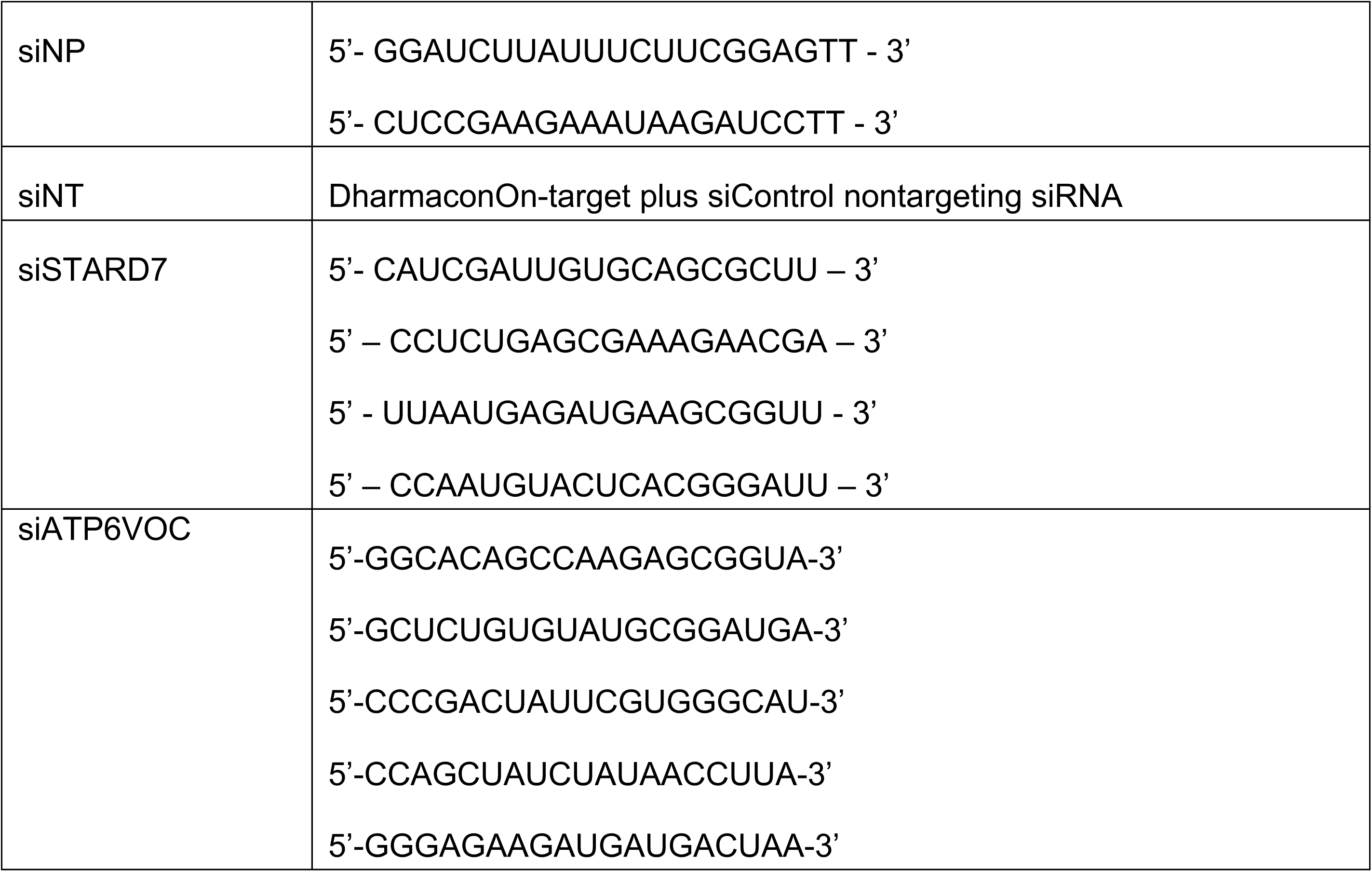

### Antibodies for immunofluorescence

Rabbit anti-STARD7 polyclonal antibodies (ab221569, Abcam), mouse monoclonal anti-NP HT103 (Mount Sinai, in-house antibody), influenza A M1 monoclonal antibody (MA1-80736, Thermo Fisher Scientific), rabbit anti-PA (GTX11899, Gene Tex), rabbit anti-PB1 (GTX125923, Gene Tex), rabbit anti-PB2 (GTX125926, Gene Tex), RanBP1 (ab97659, Abcam) and Alexa Fluor 488-conjugated and 568-conjugated goat anti-mouse secondary antibody (A-11001 and A-11004, Thermo Fisher Scientific).

### High content imaging

A549, MEF, and HTBE cells were seeded in 384-well plates at a density of 5000 cells/well and incubated overnight in serum-free media pre-spotted with compounds. The following day, cells were infected with influenza virus at the indicated MOIs and at 48hs post-infection, cells were fixed with 10% PFA for 30 min and permeabilized with 0.5% Triton X-100 for 10 min. Blocking was performed with 3% BSA for 1h, followed by incubation for 2 hs at room temperature with primary anti -NP antibody at 1:3000 dilution. After three phosphate-buffered saline (PBS) washes, the cells were incubated with an Alexa Fluor 488-conjugated anti-mouse secondary antibody at a 1:1000 dilution. Following three additional PBS washes, plates were stained with DAPI and imaged using a high content imaging system IXMC.

### Subcellular trafficking and localization studies

NP, PA, PB1, PB2, PA, RanBP1 & STARD7 localisation (Figure S2A): A549 cells were seeded in glass-bottom chambered slides (Nunc) and incubated overnight in 10% FBS DMEM media at 37 °C and 5% CO2. Cells were washed once with 1× PBS and infected with A/WSN/33 at an MOI of 3 indicated in Figure S2A for 1h. Inoculum was removed and replaced with M4 3uM in serum-free media. At indicated time points, cells were washed twice with 1× PBS and fixed using 4% paraformaldehyde (Fisher Scientific) for 30 min at room temperature. Cells were permeabilized using 0.5% Triton X-100 for 15 min, followed by 1h of blocking with 3% BSA at room temperature. This was followed by immunolabelling with indicated primary and secondary antibodies. The slides were mounted on coverslips with Prolong anti-fade DAPI (Life Technologies) and sealed. Images were acquired using the Zeiss LSM780 confocal microscope at The Scripps Research Institute Microscopy facility and localization was assessed using Fiji software.

<u>NP and M1 protein localization (</u>Figures 2C<u>, S2B, S5A)</u>: MDCK or A549 cells were seeded on glass coverslips in a 24-well plate and grown to 80% confluence overnight at 37°C and 5% CO_2_ in DMEM supplemented with 10% FBS (Gibco, Thermofisher), 1% penicillin, streptomycin, and amphotericin B mix (PAN-Biotech). Cells were pre-treated with post-inoculation media (DMEM, 0.3% BSA, 0.1% FBS, 1% penicillin, streptomycin, and amphotericin B mix) containing either M4 (3 µM) or DMSO for 2h at 37°C. Cells were washed with 1X PBS and infected with A/WSN/33 (MOI of 3) for 1h at 4°C in the presence or absence of M4. Virus inoculum was aspirated, cells were washed with 1X PBS, and replenished with post-inoculation media containing M4 (3 µM) and TPCK-treated trypsin (1 µg/mL); thereafter cells were incubated for a total of 4, 6, 8, and 12hpi. At the indicated times post infection, cells were fixed in 100% methanol for 30min at -20°C. Cells were washed twice with 1X PBS followed by a 2h block with 5% BSA at RT. Subsequently, cells were immunostained with anti-NP mouse monoclonal (HT103) or anti-M1 mouse monoclonal (MA180736) (1:2000) for 1h at RT. Hereafter, cells were washed with 1X PBS-Tween20 and incubated with Alexa Fluor 488-conjugated anti-mouse secondary antibody (1:500) and DAPI (1:1000). The coverslips were mounted on slides with ProLong Glass antifade (P36982). Images were acquired using the Nikon Eclipse 50i fluorescence microscope running NIS-Elements software (version BR 4.20.03) and further processed using ImageJ (version 1.53t) software.

STAT1-GFP trafficking (Figure 2E): Vero E6 cells in a 24-well plated were transiently transfected with 200ng STAT1-GFP plasmid DNA using Lipofectamine™ 3000 (Invitrogen, Thermofisher) according to manufacturer’s instructions. Twenty-four hours post-transfection, cells were stimulated with interferon-β (IFNβ; 1000 U/mL; IF014, Merck) in complete media supplemented with cycloheximide (CHX; 10µg/mL; C7698, Merck) and containing either DMSO, M4 (10μM), or leptomycin B (50nM). The subcellular localization of STAT1-GFP was determined by live-cell imaging at 10min, 1h, and 5h post-stimulation. Images were acquired using the Zeiss Primovert microscope equipped with the Axiocam 208 colour camera.

### Time-of-Addition Assay

MDCK cells were plated in complete DMEM and grown to 80% confluence for 24 hours. Cells were then treated with 3µM M4 at -2, 0, 2, 4, 6, 8h pre/post infection with influenza A/WSN/33 virus (MOI 0.1). Supernatants were collected 24h post infection and viral titers were quantified by plaque assay.

### Influenza virus entry assay

Replication-incompetent, vesicular stomatitis virus (VSV) engineered to express green fluorescent protein (GFP) and firefly luciferase (FLuc) in lieu of the viral glycoprotein (G), was kindly gifted by Dr. Gert Zimmer at the Institute for Virology, Mittelhäusern, Switzerland. VSV pseudoviruses bearing the HA and NA protein of influenza A/WSN/33 virus were generated by transfecting HEK-293T cells with plasmids expressing HA and NA, and then infecting the cells with VSVΔG(FLuc) that had been complemented with VSV G. Supernantants containing VSVppHA/NA were collected and titrated based on firefly luciferase expression.

For the influenza entry assay, MDCK cells were pre-treated with DMSO, M4, or S20 for 2h before infection with VSVppHA/NA in the presence of compounds (White et al., 2015a). Luciferase activity in the cell lysates was measured at 24h post infection and expressed relative to the DMSO control.

### Resistance study

The concentration of M4 required for maximum virus inhibition (3 logs), while maintaining enough virus production for subsequent passages, was determined (2 μM M4). A549 cells were infected with WSN at an MOI of 0.01 for 24 h at 37 °C under M4 treatment. The supernatant was then collected and titered by plaque assay. If the recovered M4-treated virus did not show an increased viral titer similar to that of the DMSO treated control, the virus was passaged again by the same method. This was repeated for 10 passages. The titers of DMSO and M4 passaged viruses at each passage were recorded.

### RNA extraction and RT-qPCR

For strand-specific detection of viral RNA in cytoplasmic and nuclear fractions, primers were designed for influenza virus A segment 5 positive-sense mRNA and negative-sense vRNA, each containing an additional unrelated 18- to 20-nucleotide tag at the 5′ end to enhance specificity and differentiate between the RNA species, as described previously (Kawakami et al., 2011). Briefly, equal amounts of fractionated RNA were reverse-transcribed into cDNA using the Applied Biosystems cDNA synthesis kit. qPCR was then performed using the RT2 SYBR Green qPCR Master Mix (Applied Biosystems) with primer sets specific for the corresponding influenza virus RNA (Kawakami et al., 2011) on an ABI thermocycler. Viral RNA abundance was normalized to GAPDH for cytoplasmic fractions and U6 for nuclear fractions. Relative expression levels compared to mock-treated samples were calculated using the 2^−ΔΔCT^ formula.

### Generation of CRISPR-Cas9 STARD7 knockout (KO) cells

To generate STARD7 knockout (KO) cells using CRISPR-Cas9, we targeted the STARD7 locus with the sequence ACCCTCCAGAACCAAAAGCC and generated a full-length guide RNA (gRNA) using a gRNA synthesis kit (Thermo Fisher). The gRNA was transfected into A549-dox-Cas9 cells that had been pre-treated with doxycycline (1 μg ml–1; Clontech) for 48 hs to induce Cas9 expression. At 48hs post-transfection, cells were plated for single-cell colony formation. Colonies were screened via western blot analysis to confirm STARD7 knockout. Cell viability was assessed by Western blot for STARD7 (PA5-30772, Thermo Fisher) and β-actin (4967, Cell Signaling), and cell number quantified with DAPI staining with a high content imager.

### Streptavidin enrichment studies

For streptavidin enrichment studies, confluent A549 cells grown in a 10-cm tissue culture dish were washed twice with phenol red-free DMEM (Gibco). The cells were then treated with 20 µM M4.67 or an equivalent volume of DMSO in phenol red-free DMEM without FBS and incubated at 37 °C for 1 h. Following treatment, cells were washed twice with PBS, scraped into 1 mL PBS, and then lysed by sonication. Insoluble material was removed by centrifugation. Two 1.5-mL microcentrifuge tubes, each containing 0.5 mL of lysate (2 mg/mL), were incubated with a Click reagent mix consisting of 30 µL of 1.7 mM TBTA in tBuOH:DMSO 4:1, 10 µL of 50 mM CuSO_4_ in H_2_O, 10 µL of 50 mM TCEP in H2O, 2.5 µL of 20 mM biotin-PEG3-N_3_ at room temperature for 1 h. Samples were then precipitated in ice-cold methanol, and the resulting pellet was resuspended in 1 mL PBS containing 0.6% sodium dodecyl sulfate (SDS). The resuspended lysates were pooled into a 2-mL mixture and incubated for 24hs at 4 °C with 220 µL streptavidin beads in 10 mL PBS on a rotator. Beads were pelleted by centrifugation (2,000 x*g* for 1 min) and subjected to sequential washes: two washes with 10 mL of 0.1% SDS in PBS, two washes with 10 mL of PBS twice, and two final washes with 10 mL of water. After washing, 200 µL of SDS-PAGE sample buffer supplemented with 10% beta-mercaptoethanol was added to the beads, and the beads were boiled at 99 °C for 15 min. The resulting supernatant was analyzed via western blot using anti-biotin and anti-STARD7.

### Immunoprecipitation studies

Plasmids encoding FLAG-tagged STARD7 were obtained from Origene (RC202539). Cysteine mutants were generated using site-directed mutagenesis (NEB E0554S). For immunoprecipitation studies using FLAG-tagged transgenes, HEK293T cells (5 × 10^6^) were transfected with 2 µg of each plasmid per well of a six-well plate using 100 µL of OptiMEM medium (Gibco) containing 8 µL of FuGENE HD transfection reagent (Promega). After 24hs, the growth mediμM replaced with fresh medium, and cells were incubated for another 24 hs. At 48hs post-transfection, cells were treated with 20 µM M4.67 in serum-free DMEM for 1h. Following treatment, cells were washed twice with PBS, scraped into 250 µL of ice-cold RIPA buffer (EMD Millipore), and lysed by sonication. Insoluble material was removed by centrifugation, and protein concentration was determined via absorbance measurements. For immunoprecipitation, 1mg of total lysate in (1 mL of RIPA buffer) was incubated overnight at 4 °C with 20 µL of anti-FLAG M2 magnetic bead slurry (Sigma). Beads were washed three times with 300 µl of RIPA, and bound protein complex were eluted using 250 µg/mL of FLAG peptide (DYKDDDDK, Sino Biological) in PBS. Eluted samples were then subject to click reaction (described previously) and incubated at room temperature for 1 h. The reaction mixture was precipitated in ice-cold methanol, and the pellet was resuspended in 50µL SDS-PAGE sample buffer with 10% beta-mercaptoethanol. Immunoprecipitated proteins were separated by SDS-PAGE and analyzed using a ChemiDoc MP imager (Bio-Rad) via rhodamine fluorescence scanning and anti-FLAG immunoblotting.

To examine NP interactions with viral proteins during infection, A549 cells were seeded in 10-cm dishes, pre-treated with M4 (5 µM) or DMSO for 16 h at 37 °C, and infected with influenza A/WSN/33 at an MOI of 3 for 1 h at 4 °C. Following virus adsorption, inoculum was removed and cells were incubated in infection medium containing M4 (5 µM) or DMSO for 8 h at 37 °C.

Cells were washed with ice-cold PBS and lysed in ice-cold Pierce IP lysis buffer supplemented with protease and phosphatase inhibitors. Lysates were clarified by centrifugation at 5,000 × g for 20 min at 4 °C. A fraction of each lysate was retained as input. Remaining lysates were incubated overnight at 4 °C with Protein G magnetic beads pre-bound to anti-influenza A nucleoprotein antibody (Genetex, GTX125989).

Beads were washed three times with Pierce IP lysis buffer, and bound proteins were eluted in 1× Laemmli buffer by boiling at 95 °C. Input and immunoprecipitated samples were resolved by SDS-PAGE and transferred to PVDF membranes (Immobilon-P, Merck). Membranes were blocked in 3% BSA in TBST and probed with antibodies against influenza A NP (Genetex, GTX125989), M1 (Invitrogen, MA1-80736), NS2/NEP (Genetex, GTX125952), and GAPDH (Genetex, GTX627408), followed by HP-conjugated secondary antibodies. Proteins were detected using enhanced chemiluminescence (ECL Prime, Cytiva).

### Animal Studies

For the *in vivo* studies, 11-12-week-old male or female C57BL/6 mice (Jackson Laboratories, Bar Harbor, ME, USA) were randomly assigned to experimental groups. All procedures and handling adhered to the current standards specified in the Guide for the Care and Use of Laboratory Animals, and all experiments were conducted under an IACUC-approved protocol (24-0006-1; 07/01/2022-06/30/2032; IACUC at The Scripps Research Institute) in a USDA-registered research facility. The mice were bred and housed in a BSL2 barrier facility following institutional guidelines. In these studies, mice were divided into experimental groups of 3 and received different treatments. Vehicle (0.2% methylcellulose and Tween 80) and baloxivir was administered orally once 12hs prior to infection, while M4 was given intraperitoneally twice daily, starting 12 hours before infection and continuing until day 4 post-infection. For virus challenges, mice were anesthetized using the open-drop method of isoflurane exposure and then received an intranasal administration of 300 PFU of influenza A/WSN/33 (H1N1) in a 30 μL volume. On day 5 post-infection, 5 mice were euthanized, and their lungs were collected, homogenized in 1 mL of DMEM, and stored at −80 °C until titration was performed using the standard plaque assay in MDCK cells. The remaining 3 mice were monitored for body weight for 5 days.

## Acknowledgements

This work was partly funded by grant W81XWH-20-1-0270 from the US Department of Defense to AG-S, by CRIPT (Center for Research on Influenza Pathogenesis and Transmission), a NIAID funded Center of Excellence for Influenza Research and Response (CEIRR, contract# 75N93021C00014) and by ARPA-H agreement AY2AX000073-01 to AG-S

## Disclosures

The A.G.-S. laboratory has received research support from Avimex, Dynavax, Pharmamar, 7Hills Pharma, ImmunityBio and Accurius, outside of the reported work. A.G.-S. has consulting agreements for the following companies involving cash and/or stock: Castlevax, Amovir, Vivaldi Biosciences, Contrafect, 7Hills Pharma, Avimex, Pagoda, Accurius, Esperovax, Applied Biological Laboratories, Pharmamar, CureLab Oncology, CureLab Veterinary, Synairgen, Paratus, Pfizer, Virofend and Prosetta, outside of the reported work. A.G.-S. has been an invited speaker in meeting events organized by Seqirus, Janssen, Abbott, Astrazeneca and Novavax. A.G.-S. is inventor on patents and patent applications on the use of antivirals and vaccines for the treatment and prevention of virus infections and cancer, owned by the Icahn School of Medicine at Mount Sinai, New York, outside of the reported work.

## Supporting information

**Figure S1A:**
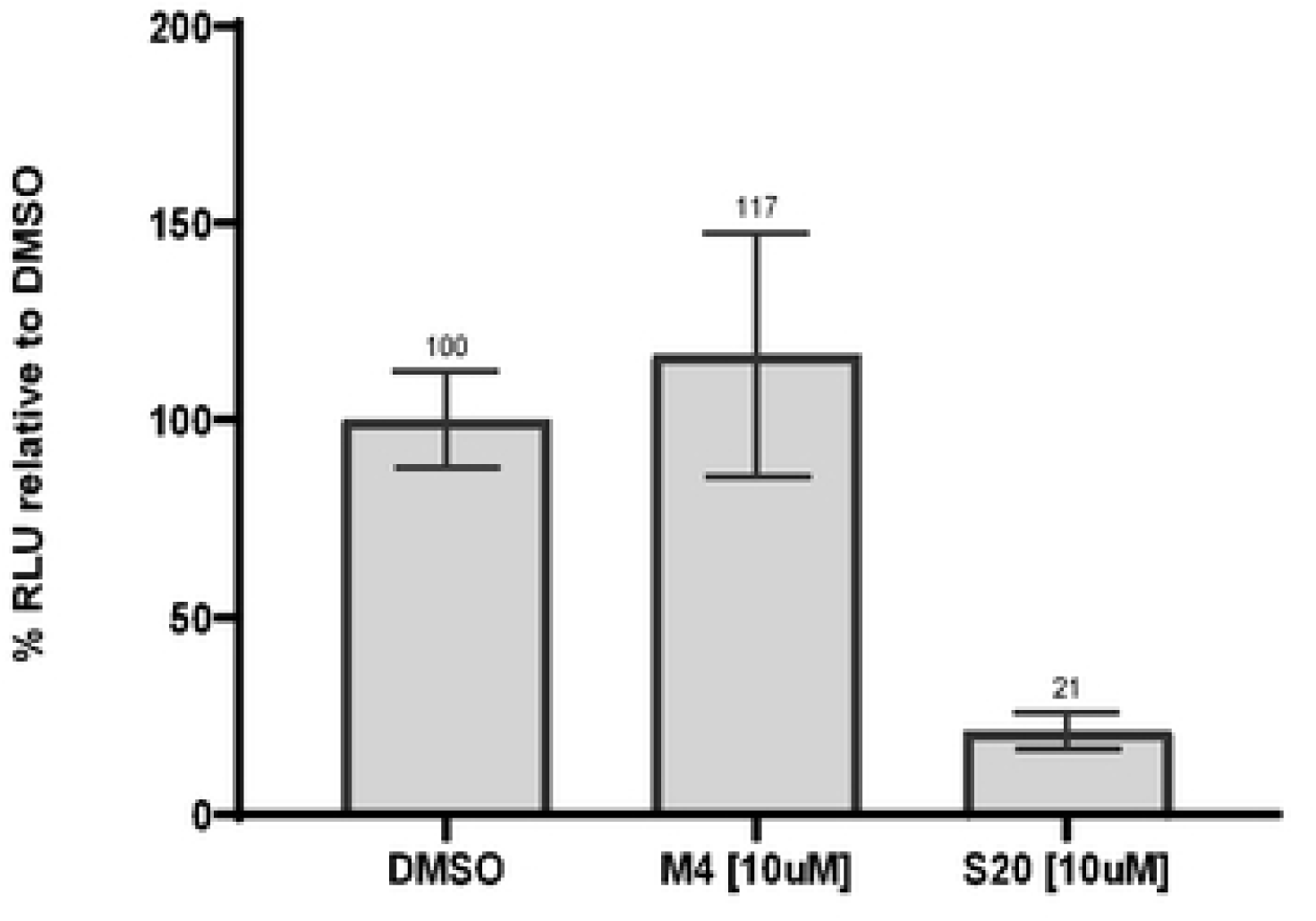
MDCK cells were infected with influenza HA/NA-pseudotyped VSV particles in the presence of DMSO, M4, or the HA fusion inhibitor S20. Luciferase activity was measured at 24 hours post-infection to assess entry efficiency. The experiment was performed in triplicate and the means +/- standard deviation are shown.

**Figure S2A:**
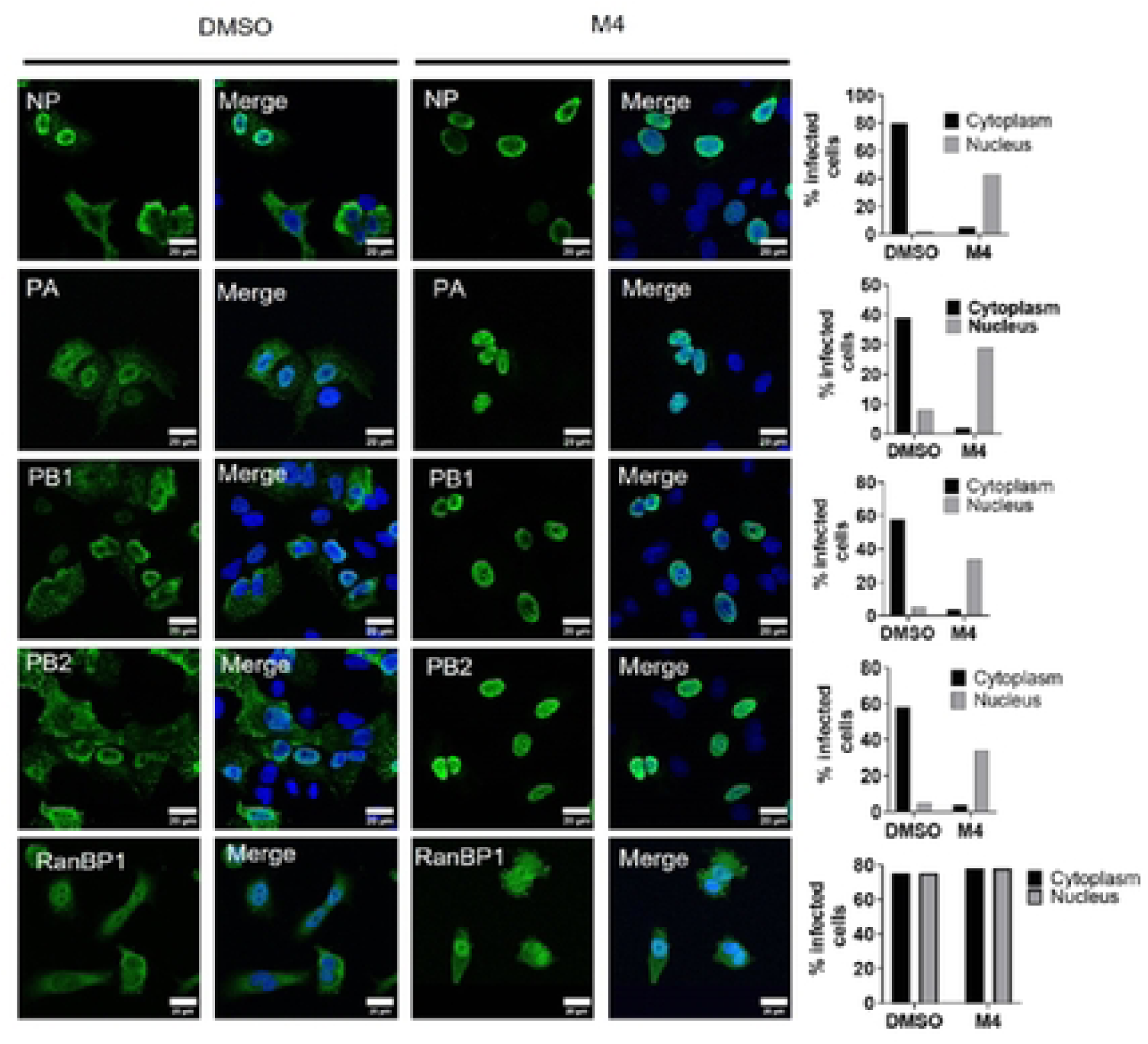
A549 cells were pre-treated for 2h with 3μM M4 followed by infection with A/WSN/33 at 3 MOI for 24h. Cells were then fixed, labeled with anti NP, PA, PB1, PB2, or RanGP1 antibodies and analyzed for immunofluorescence. Representative images are shown. Quantification in the last panel of each row was done with Image J.

**Figure S2B:**
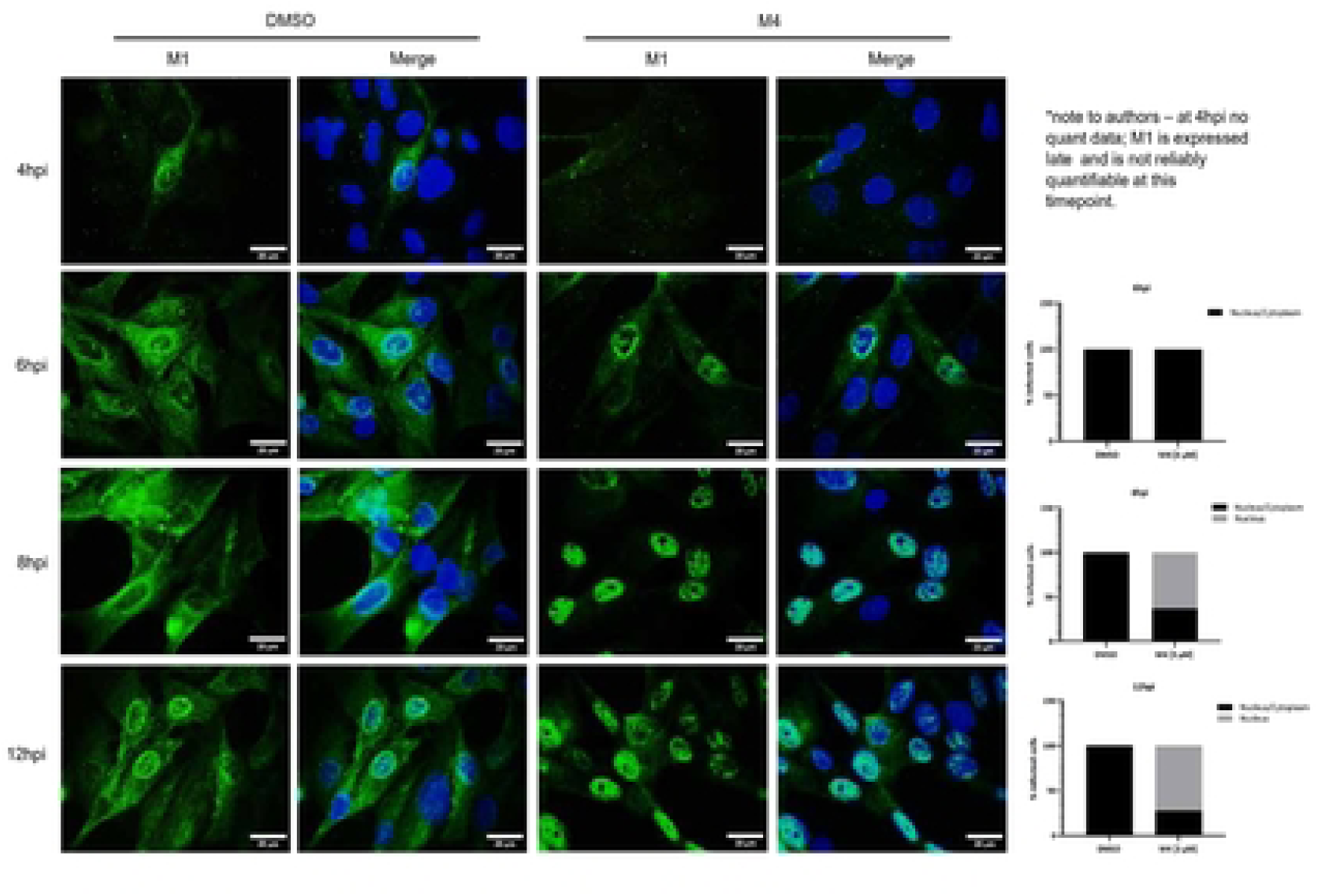
Subcellular localisation of influenza A virus M1 protein at indicated times post-infection in M4 treated cells. MDCK cells treated with M4 (3 µM) and infected with influenza A/WSN/33 virus (MOI 3) were immunolabeled with anti-M1 antibodies at 4, 6, 8, and 12hpi to capture the cellular trafficking of M1 proteins. Scale bar = *20 µm*. Five fields of view were randomly selected for M4 and DMSO, respectively. The localisation of M1 was recorded as present either in the “nucleus” or “nucleus/cytoplasm” and graphically displayed as a percentage (%).

**Figure S2C:**
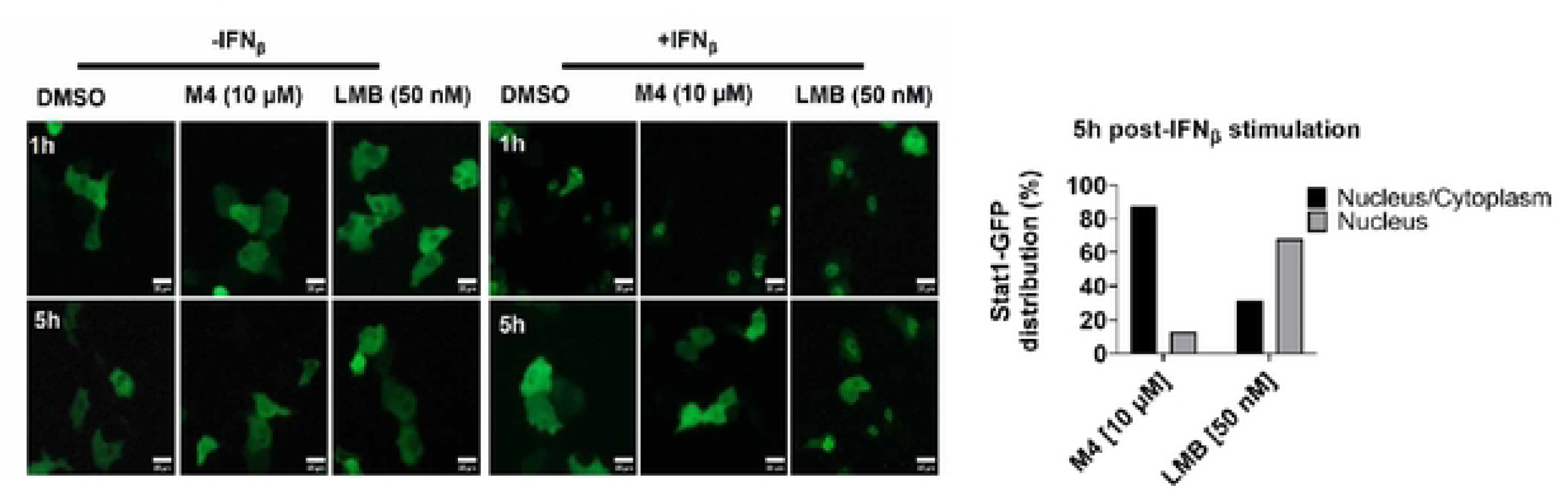
Vero E6 cells transiently expressing STAT1-GFP are shown in the presence of DMSO, M4, or Leptomycin B (LMB). Cells were either unstimulated or stimulated with IFNβ, and live-cell images of the subcellular localization of STAT1-GFP were collected by fluorescent microscopy at the indicated time points (1 h, and 5 h). Scale bar = 25 µm. For quantification, 29 fields of view were randomly selected for M4 and LMB, respectively. The localization of STAT1-GFP was recorded as present either in the “nucleus” or “nucleus/cytoplasm”, and the data are displayed as a percentage (%).

**Figure S2D:**
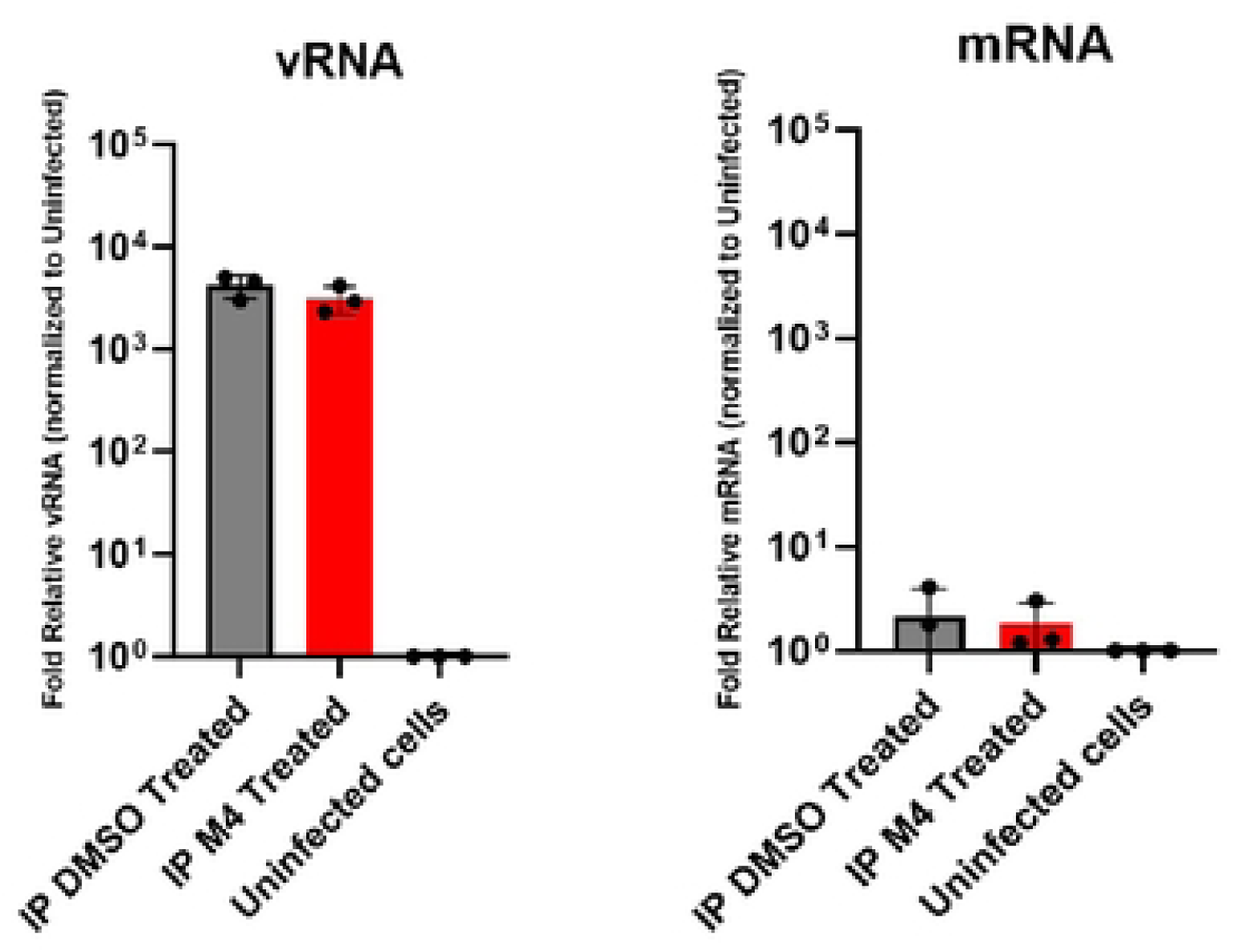
vRNA and mRNA associated with NP immunoprecipitates were quantified by RT-qPCR. A549 cells were infected with influenza A virus (WSN) and treated with either DMSO or M4 (1 µM). NP–vRNP complexes were isolated by immunoprecipitation, and viral RNA levels were measured using segment-specific primers. Ct values were converted to relative quantities using the 2^–ΔCt method with uninfected cells serving as the reference. Data represent fold relative vRNA (left) and viral mRNA (right) associated with NP immunoprecipitates and are presented as the mean ± SEM from thee biological replicates.

**Figure S3A:**
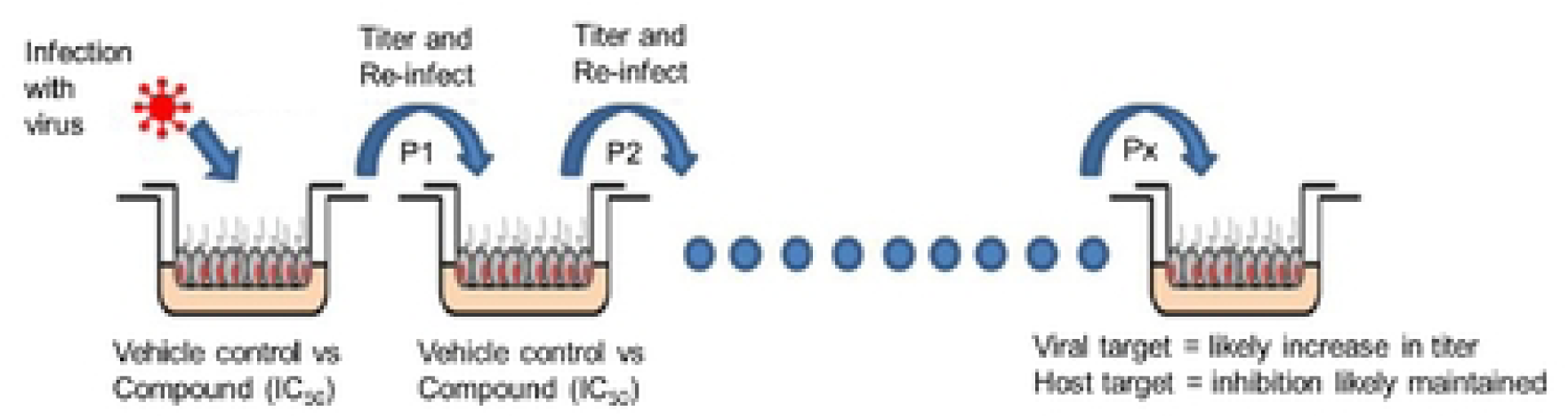
Schematic of the serial passaging workflow.

**Figure S3B:**
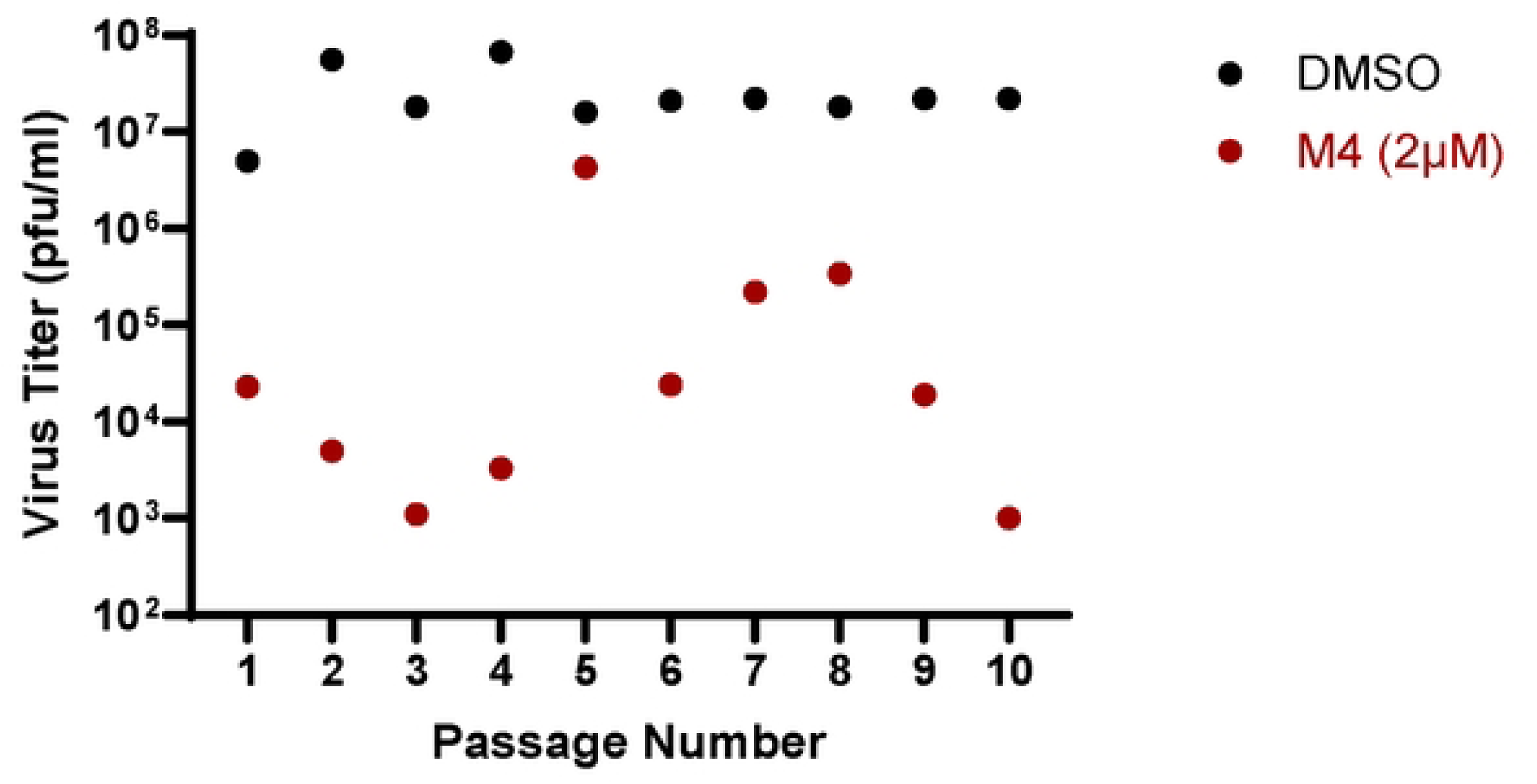
M4 response measurements collected across passages. Data points show values obtained at each passage for M4 (2 μM), plotted as individual measurements

**Figure S4A:**
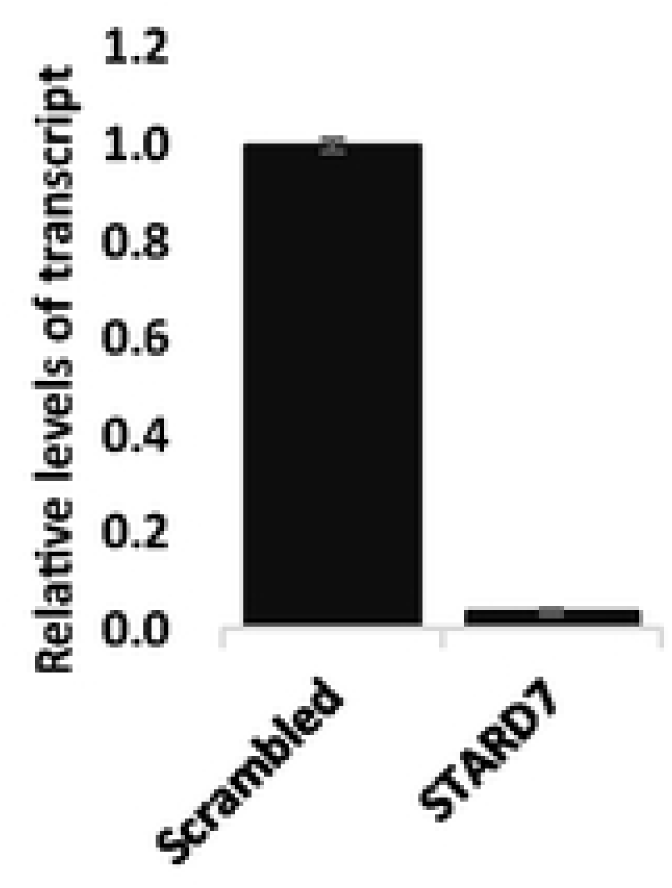
STARD7 knockdown efficiency by qPCR. A549 cells were transfected with either non-targeting (NT) siRNA or STARD7 siRNA, and mRNA levels were quantified by qPCR using STARD7-specific primers. Data are normalized to a GAPDH housekeeping gene and presented as fold change relative to NT siRNA. Statistical significance was determined with a student t-test. ****p<0.01

**Figure S5A:**
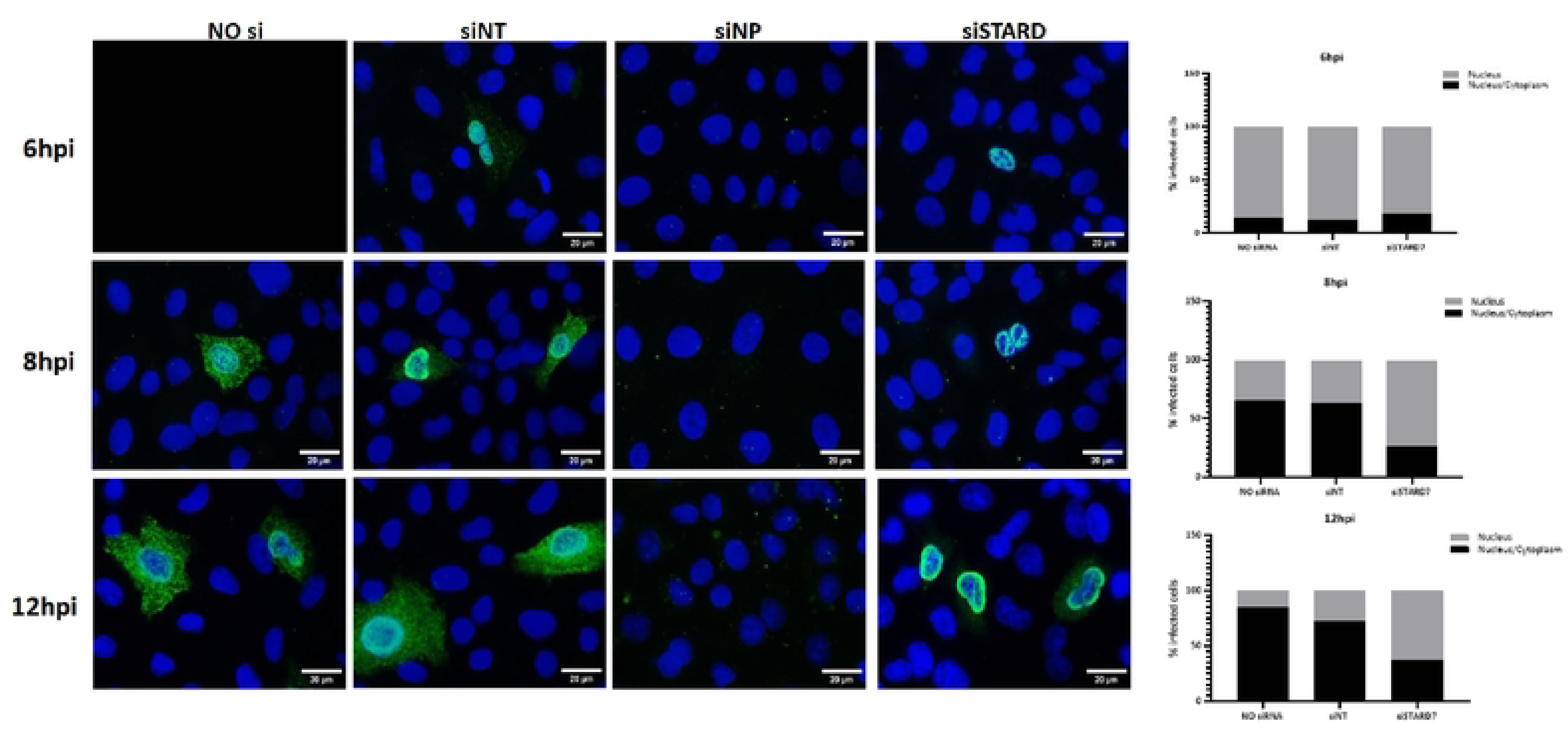
Subcellular trafficking of influenza A virus NP in STARD7 knockdown cells. A549 cells were transfected with no siRNA, non-targeting control siRNA, STARD7-targeting siRNA, or NP-targeting siRNA for 48hs, followed by infection with influenza A/WSN/33 virus. Subcellular localisation of NP was captured by indirect immunofluorescence microscopy at 6-, 8-, and 12-hours post-infection. Scale bar = *20 µm.* An average of 70 cells were quantified per condition at each timepoint. NP localisation was scored as present either “in the nucleus” or “both in the nucleus and cytoplasm (nuclear/cyto)” and the ratios are graphically shown as a percentage (%).

**Figure S6A:**
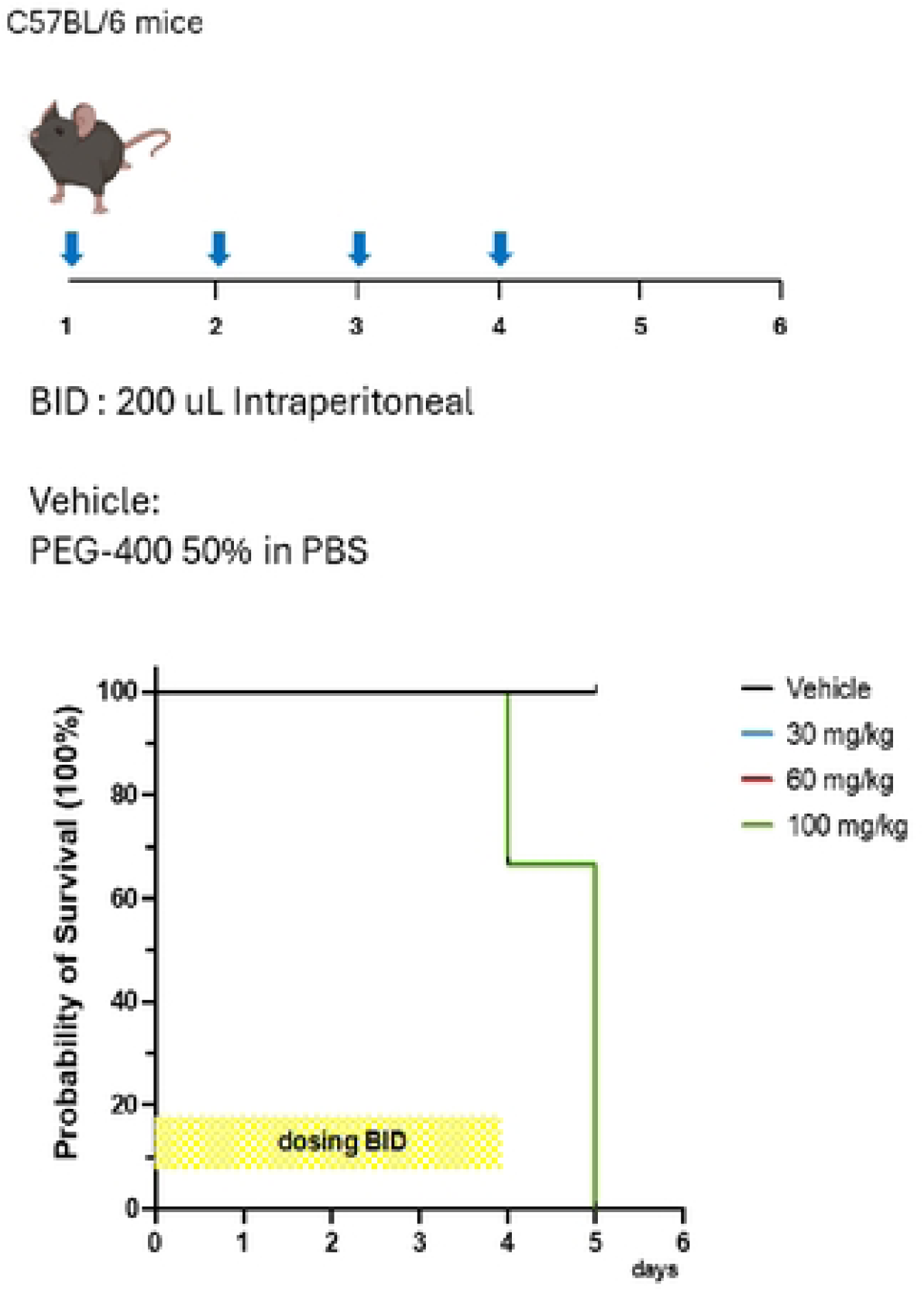
Schematic of the dosing regimen for the M4 maximum tolerated dose study. Mice were administered M4 intraperitoneally (IP) twice daily (BID) for six consecutive days at doses of 30 mg/kg, 60 mg/kg, or 100 mg/kg. The vehicle was 50% PEG-400 in PBS.

**Figure S6B.**
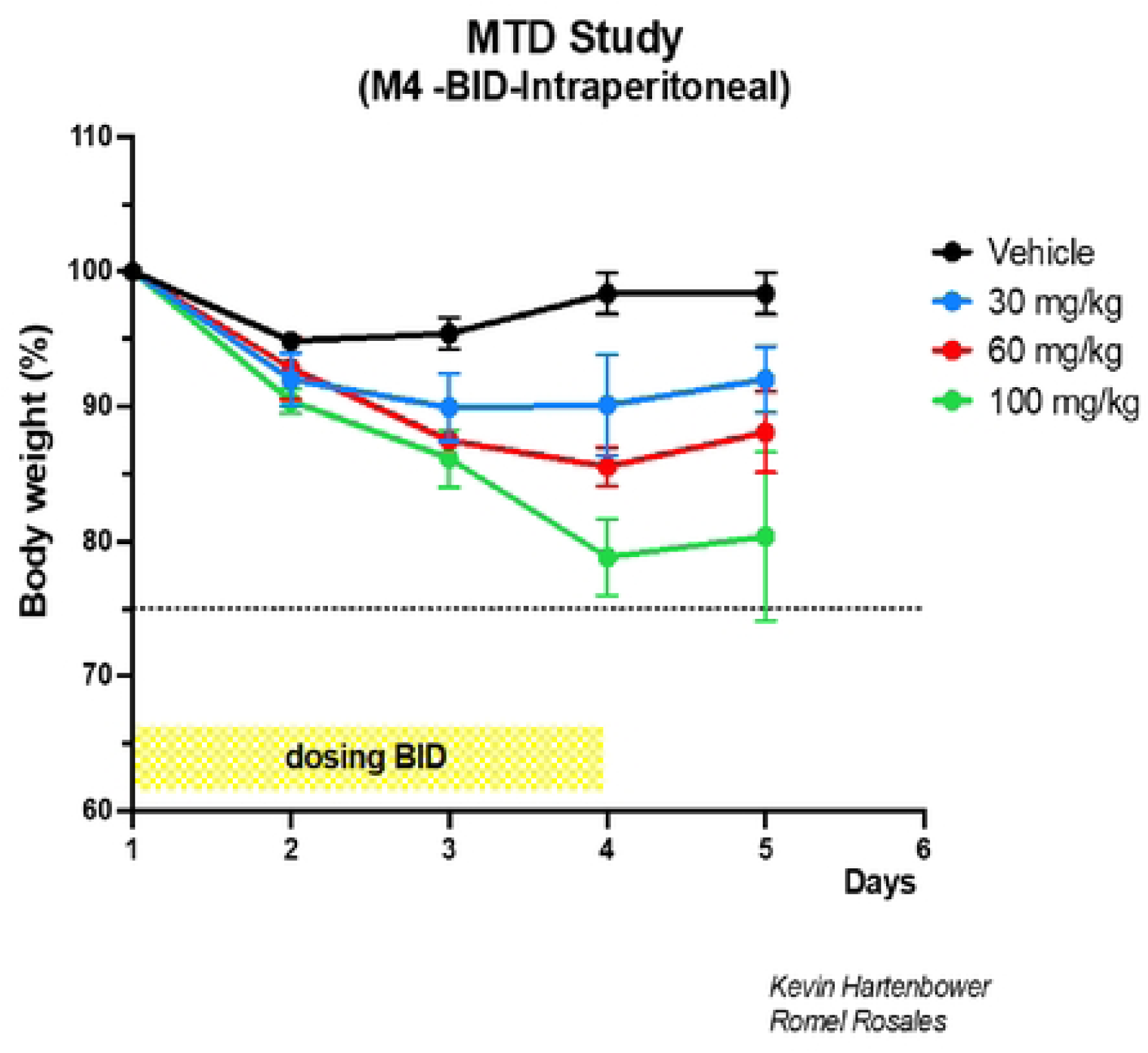
Body weight was recorded daily and expressed as a percentage of baseline (Day 0). Mice receiving 100 mg/kg exhibited weight loss exceeding 10% of starting body weight by Day 3, triggering humane euthanasia per IACUC guidelines.

**Figure S6C** Kaplan–Meier survival analysis over the 6-day treatment period.

**Table S1:**
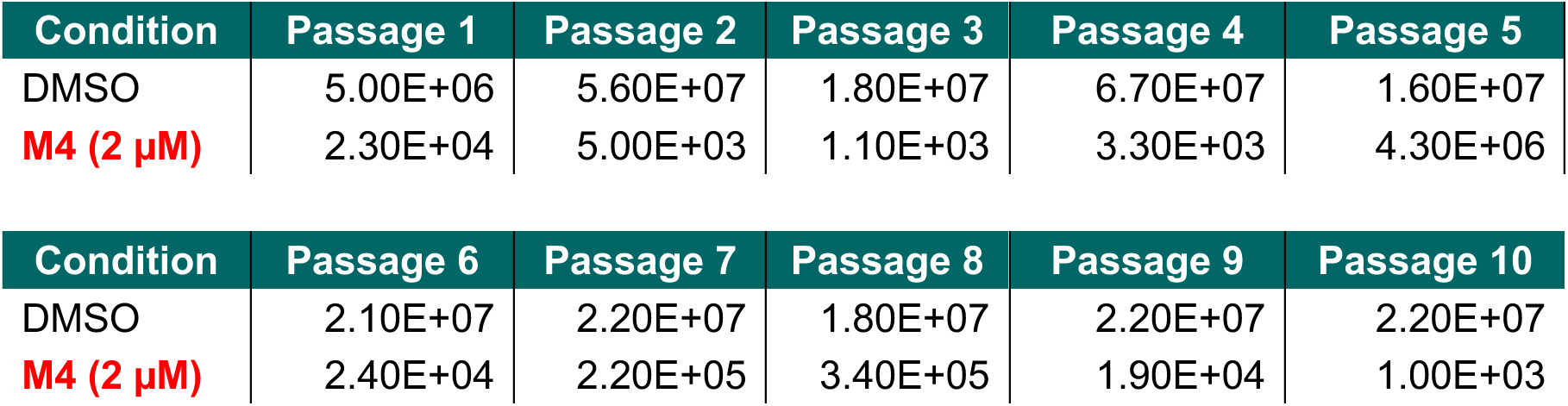
M4 treatment fails to select for resistant virus: Influenza A/WSN/33 virus was serially passaged ten times in the presence of DMSO or M4 (2 µM). Black points represent DMSO-treated controls, and red points represent M4-treated cultures. Individual data points correspond to titers measured at each passage. Corresponding quantitative titers for each passage are provided in Table S1.

**Table S2A:** Top enriched proteins as high priority targets

| Symbol | Gene ID | 10 μM log2 FC | 1 μM log2 FC |
| --- | --- | --- | --- |
| STARD7 | StAR-related lipid transfer protein 7, mitochondrial | 9.77 | 9.48 |
| FNBP1L | Formin-binding protein 1 | 9.19 | 8.73 |
| ADO | 2-aminoethanethiol dioxygenase | 8.65 | 7.69 |
| RNASEH2A | Ribonuclease H2 subunit A | 8.34 | 7.66 |
| LRRFIP2 | Leucine-rich repeat flightless-interacting protein 2 | 8.09 | 7.2 |
| HBQ1 | Hemoglobin subunit theta-1 | 8.07 | 8.08 |
| CNN2 | Calponin-2 | 7.99 | 7.12 |
| RMI2 | RecQ-mediated genome instability protein 2 | 7.49 | 7.27 |
| C17orf75 | Protein Njmu-R1 | 7.34 | 7.36 |
| BTAF1 | TATA-binding protein-associated factor 172 | 7.29 | 7.42 |

**Table S2B:** b and y ion designations from the modified tryptic STARD7 peptide containing C302, as shown in Figure 3G.

| # | b | b<sup>++</sup> | b<sup>0</sup> | b<sup>0++</sup> | Seq. | y | y<sup>++</sup> | y<sup>*</sup> | y<sup>*++</sup> | y<sup>0</sup> | y<sup>0++</sup> | # |
| --- | --- | --- | --- | --- | --- | --- | --- | --- | --- | --- | --- | --- |
| 1 | 164.0706 | 82.5389 |  |  | Y |  |  |  |  |  |  | 17 |
| 2 | 656.1446 | 328.5759 |  |  | C | 2236.8660 | 1118.9367 | 2219.8395 | 1110.4234 | 2218.8555 | 1109.9314 | 16 |
| 3 | 755.2130 | 378.1101 |  |  | V | 1744.7921 | 872.8997 | 1727.7655 | 864.3864 | 1726.7815 | 863.8944 | 15 |
| 4 | 842.2450 | 421.6261 | 824.2345 | 412.6209 | S | 1645.7237 | 823.3655 | 1628.6971 | 814.8522 | 1627.7131 | 814.3602 | 14 |
| 5 | 1028.3243 | 514.6658 | 1010.3138 | 505.6605 | W | 1558.6916 | 779.8495 | 1541.6651 | 771.3362 | 1540.6811 | 770.8442 | 13 |
| 6 | 1175.3597 | 588.1835 | 1157.3492 | 579.1782 | M | 1372.6123 | 686.8098 | 1355.5858 | 678.2965 | 1354.6018 | 677.8045 | 12 |
| 7 | 1274.4281 | 637.7177 | 1256.4176 | 628.7124 | V | 1225.5769 | 613.2921 | 1208.5504 | 604.7788 | 1207.5664 | 604.2868 | 11 |
| 8 | 1361.4602 | 681.2337 | 1343.4496 | 672.2284 | S | 1126.5085 | 563.7579 | 1109.4820 | 555.2446 | 1108.4979 | 554.7526 | 10 |
| 9 | 1448.4922 | 724.7497 | 1430.4816 | 715.7445 | S | 1039.4765 | 520.2419 | 1022.4499 | 511.7286 | 1021.4659 | 511.2366 | 9 |
| 10 | 1505.5137 | 753.2605 | 1487.5031 | 744.2552 | G | 952.4444 | 476.7259 | 935.4179 | 468.2126 | 934.4339 | 467.7206 | 8 |
| 11 | 1652.5491 | 826.7782 | 1634.5385 | 817.7729 | M | 895.4230 | 448.2151 | 878.3964 | 439.7019 | 877.4124 | 439.2098 | 7 |
| 12 | 1749.6018 | 875.3046 | 1731.5913 | 866.2993 | P | 748.3876 | 374.6974 | 731.3610 | 366.1842 | 730.3770 | 365.6921 | 6 |
| 13 | 1864.6288 | 932.8180 | 1846.6182 | 923.8127 | D | 651.3348 | 326.1710 | 634.3083 | 317.6578 | 633.3243 | 317.1658 | 5 |
| 14 | 2011.6972 | 1006.3522 | 1993.6866 | 997.3469 | F | 536.3079 | 268.6576 | 519.2813 | 260.1443 | 518.2973 | 259.6523 | 4 |
| 15 | 2124.7812 | 1062.8943 | 2106.7707 | 1053.8890 | L | 389.2395 | 195.1234 | 372.2129 | 186.6101 | 371.2289 | 186.1181 | 3 |
| 16 | 2253.8238 | 1127.4156 | 2235.8133 | 1118.4103 | E | 276.1554 | 138.5813 | 259.1288 | 130.0681 | 258.1448 | 129.5761 | 2 |
| 17 |  |  |  |  | K | 147.1128 | 74.0600 | 130.0863 | 65.5468 |  |  | 1 |

## Notes

### Competing Interest Statement

The authors have declared no competing interest.

